# Correlative X-ray imaging and fluorescence microscopy

**DOI:** 10.64898/2026.06.04.730059

**Authors:** Mangalika Sinha, Boram Yu, Rita Mendes da Silva, Ulrike Rölleke, Peter Luley, Malte Tiburcy, Wolfram-Hubertus Zimmermann, Manfred Burghammer, Sarah Köster

## Abstract

Imaging the structural organization inside cells in their native state is essential for understanding how the arrangement and interactions among molecular components give rise to biological function. Fluorescence microscopy is one of the pivotal techniques that provides molecular specificity for imaging in real space, however, the technique is limited to labeled components. X-rays, on the contrary, are sensitive to electron density contrast and therefore to label-free samples, and probe structure in reciprocal space. In particular, scanning small-angle X-ray scattering (SAXS) combines information from real and reciprocal space and enables access to intact cells, owing to the high penetration power of the X-rays. Combining both imaging modalities in a synergistic manner promises powerful tools for cellular imaging, but remains challenging, because of the differing requirements the complementary methods introduce. Here we present a correlative imaging platform that integrates a modular, compact and beamline-compatible fluorescence microscope with scanning SAXS, to enable fast sequential imaging of the identical cellular regions. We developed a dedicated microfluidics flow chamber enabling measurements under hydrated, near-native conditions. We demonstrate the utility of our methodology by investigating two different relevant cellular components, i.e., thick keratin bundles in epithelial cells that contribute to cell mechanics, and force-generating actomyosin in cardiomyocytes. Employing adapted data analysis methods, we find a good agreement between the fluorescence-derived and the SAXS-derived orientation maps. This result demonstrates that the label-free approach with SAXS captures cytoskeletal organization throughout the cell, and can be directly linked to specific molecular information provided by the complementary fluorescence imaging, in a physiologically relevant cellular environment. Our work establishes a general strategy for multimodal imaging of cellular architecture and opens ways to investigate living cells under the influence of drugs and chemical manipulation experiments.

## Introduction

Biological function is to a great extent determined by the structural and dynamic properties of the cellular components, including the cytoskeleton, membranes and organelles, such as the nucleus. For this reason, imaging of intracellular structures at high spatial resolution has become an invaluable tool in the life sciences. In particular, correlative imaging that combines two or more different methods reveals complementary information about the system under investigation. Here we combine visible light fluorescence microscopy and X-ray scattering to collect protein-specific structural data from biological cells. In fluorescence microscopy, targeted labeling of specific proteins of interest offers the possibility to selectively visualize these components inside cells. X-ray scattering, by contrast, is a label-free technique, which records scattered photons from all matter within the beam and thereby images electron density contrast directly. X-rays have a small wavelength, enabling high resolution, and a high penetration power, allowing us to image “thick” samples like whole cells.

Among the various X-ray imaging techniques that have been developed in recent decades [1–3], scanning small-angle X-ray scattering (SAXS) is an advancement of solution SAXS with a focused X-ray beam being scanned with respect to the sample plane; a spatially resolved scattering signal is thus recorded from each scan position. Therefore, scanning SAXS in an elegant manner combines real space imaging and reciprocal space structural information. Importantly, the size of the scatterers, i.e., the intracellular components, and the beam diameter are of the same order – tens to hundreds of nanometers. Thus, “classical” analysis approaches known from solution SAXS fail and innovative data analysis needs to be developed and applied [4]. When first developed, scanning SAXS has been applied to comparatively radiation-resistant biomaterials such as wood [5], bone [6–8], teeth [9, 10], and tendons [11]. More recently, structures in freeze-dried biological cells [12] including keratin bundles [13, 14], actin stereocilia [15], and the DNA in the cell nucleus [16] have been studied. The advantage of studying dry samples lies in the improved electron density contrast between the surrounding air and the cellular material. It is therefore relatively straight-forward to reveal quantitative information on structural parameters such as bundle and filament orientation and alignment [13] and filament-packing geometries within a bundle [14]. However, fixed-hydrated or even living samples are of great interest as they come much closer to the native situation of a biological cell [17, 18]. Beside the diminished electron density contrast, the increased sensitivity to radiation damage of such samples poses great challenges.

To provide such a near-native, close-to-physiological environment, dedicated microfluidic chambers have been developed and applied [19–22]. Such flow chambers provide a number of advantages, as the cells are constantly provided with nutrients, waste is transported off, the sample is slightly cooled by the liquid flow, and, importantly, the generation of gas bubbles by the radiation that would prevent further measurements, is greatly diminished. Using such chambers and thereby enabling long total experiment times, recently, fast continuous scanning of large sample areas and, therefore, high numbers of cells was achieved [20, 23]. While microfluidic chambers for sole microscopy applications fabricated by soft lithography [24] are a standard tool in many labs now, the fabrication of a sample chamber that is equally well suitable for microscopy and for X-ray imaging remains challenging. This challenge was also identified in a recent study that applied the combination of X-ray imaging and super-resolution fluorescence microscopy to biological cell samples [25, 26]: due to the lack of compatible sample chambers, the cells were studied in freeze-dried state. Nonetheless, the study impressively showed the great advantage of combining different imaging modalities in a complementary manner.

Here, we present the design, construction and application of a compact, high-quality fluorescence microscope that is integrated in a synchrotron beamline in a straight-forward manner. We combine the microscope with a dedicated sample environment, i.e., a microfluidic chamber that fulfills all necessary requirements: it is compatible with scanning SAXS and transparent to X-rays, it is compatible with fluorescence microscopy and moderate working distance objectives, it includes a thin water layer only, thus providing good X-ray contrast, and it enables continuous flow to diminish X-radiation damage and gas bubble formation. We apply the novel development to two cellular systems with particularly well-ordered structures: first, pronounced keratin bundles in SW-13 cells and second, force generating actomyosin in iPS cardiomyocytes. Our approach enables a smooth workflow that was previously missing, including a compact design suitable for various beamlines, fast switching between the imaging modalities, the possibility for extended investigation of biological cells in a near-native, aqueous environment and the applicability to different (biological and non-biological) sample types. In this work, we focus on quantifying ordered structures in both fluorescence microscopy and scanning SAXS data, and the orientation maps derived from both modalities show great agreement. Thus, our setup demonstrates the benefit of multimodal imaging of biological systems.

## Results

### Integrated fluorescence microscope setup

Combined X-ray imaging and visible light fluorescence microscopy promises to provide highly complementary information from biological samples; while scanning SAXS probes structural parameters like alignment and orientation of subcellular components, in a spatially-resolved manner, fluorescence microscopy highlights the specifically labeled components and locates them within the context of the cell. To combine both methods, we develop a compact epi-fluorescence microscope specifically designed for integration into a synchrotron beamline. Fig. 1a shows a schematic of the optical path of the microscope, which comprises a light source and light guide, an aspheric collimation lens, an excitation filter, a dichroic mirror, a high-NA objective, an emission filter, a tube lens and a sensitive camera, arranged to minimize footprint while maintaining excellent optical performance. Here, a GFP filter set is used, however, it can be changed to different wavelengths in a straight-forward manner. The specifications of all the optical components used in the set-up are provided (Supplementary Table 1). We employ a 20*×* , NA 0.7 objective with a working distance of 1 mm, as an optimal compromise between sufficiently high lateral resolution, a large field of view, and geometrical compatibility with the microfluidic chamber in the beamline set-up. Typically, higher-magnification objectives have much shorter working distances on the order of 0.1-0.3 mm, which are incompatible with the microfluidic chamber geometry. Specifically, we use fragile Si_3_N_4_ membranes of 1 *µ*m thickness of the window material as it is compatible with SAXS, microscopy and cell culture. This prohibits the use of immersion objectives that require close proximity substrate and risk damaging the membrane. The microscope is mounted on a motorized linear stage that translates the optical axis in and out of the X-ray path with a reproducibility down to 1 *µ*m, enabling fluorescence imaging without perturbing the specimen position relative to the nanofocused X-ray beam. Fig. 1b shows a photograph of the microscope within the X-ray path and in relation to the sample chamber. The main components are colored and explained in the legend.

**Figure 1:**
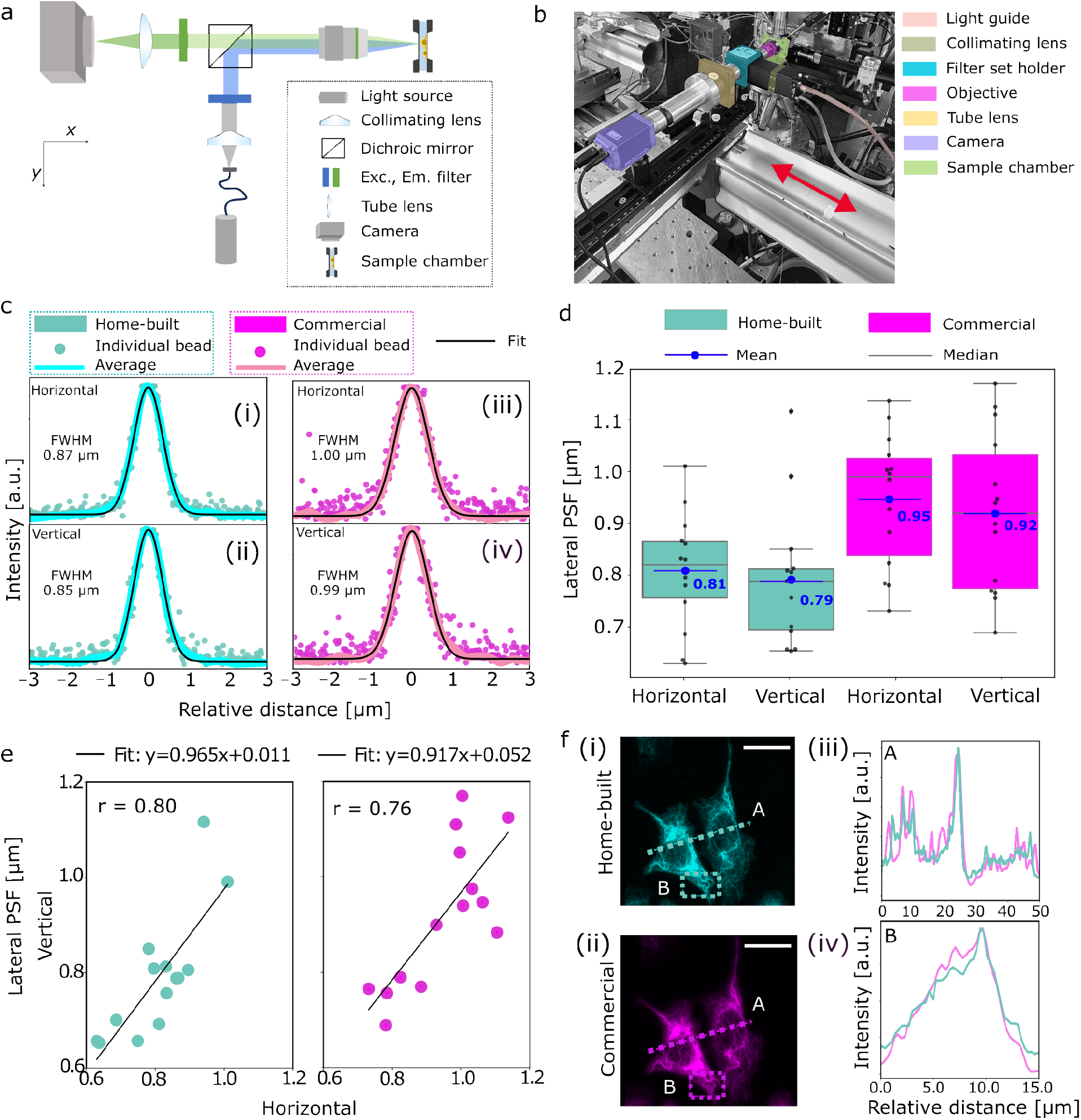
Design and implementation of a fluorescence microscope for integration in a synchrotron beam-line. (a) Schematic of the home-built epi-fluorescence microscope showing the optical layout optimized for integration in the beamline. (b) Photograph of the microscope installed at the beamline (ESRF, ID13). The relevant optical components are highlighted in different colors. In this geometry the sample stays in the X-ray beam path; fluorescence imaging is performed sequentially to the X-ray measurements by shuttling the microscope in and out of X-ray beam path, as indicated by the red arrow. (c) Evaluation of the lateral PSF of the microscope using 0.1 *µ*m-diameter fluorescent beads suspended in ultrapure water and deposited between two Si_3_N_4_ windows arranged in analogy to the actual measurements. (i,ii) Show the individual horizontal and vertical line intensity profiles of N = 14 beads (cyan, scatter) along with the averaged intensity profile (cyan, solid line) and the corresponding Gaussian fit (black, solid line). (iii, iv) Show the the corresponding measurements for a commercial epi-fluorescence microscope (shown in magenta). Identical objectives and filter sets are used in both configurations. The FWHM denotes the lateral point spread function of the averaged horizontal and vertical profiles. (d) Quantitative comparison of the lateral point spread function between the home-built beamline microscope and the commercial system. The FWHM from the Gaussian fits of the individual horizontal and vertical line profiles of each of the N= 14 beads for each case are plotted as single data points (black). The box plots indicate the range of values between 25th percentile and 75th percentile with the median (gray) and the mean (blue) of the distributions. (e) Correlation of bead FWHM in the horizontal and vertical axes in the focal plane, illustrating the degree of astigmatism. Each point represents an individual bead; the black lines show linear fits. (f) Images of keratin bundles in cells acquired from the same region on a sample using the home-built and a commercial microscope (scale bars: 20 *µ*m). The normalized intensity profiles for **A** (dotted line) and **B** (dotted box) agree well.

To assess the quality of the imaging achieved by our home-built fluorescence microscope, we benchmark it against a commercial research microscope equipped with the same objective and filter set. We determine the lateral point spread function (PSF) by preparing samples of 0.1 *µ*m diameter yellow-green fluorescent beads dissolved in ultrapure water at a final concentration of 0.1 *µ*g/mL, on plasma-activated Si_3_N_4_ membrane windows. The beads are allowed to adhere during a one hour incubation period. After incubation, the bead-coated window is integrated within the microfluidic chamber specifically designed for this study and the beads in aqueous environment are imaged. Images of individual beads are analyzed by extracting horizontal and vertical intensity profiles through the centroids. The intensity profiles from N = 14 individual beads are first smoothed, normalized, aligned, linearly interpolated and subsequently averaged to obtain representative average bead profiles for the horizontal (H) and vertical (V) directions. Gaussian functions are then fitted to the averaged bead profiles and the lateral PSF is determined as the full width at half maximum (FWHM) (Fig. 1c). The home-built microscope yields results for the FWHM of 0.87 *±*0.11 *µ*m (H) and 0.85 *±*0.14 *µ*m (V), indicating a sub-micrometer PSF with minimal astigmatism (Fig. 1c(i,ii)). Measurements with the commercial microscope yield mean FWHM values of 1.00 *±*0.13 *µ*m (horizontal) and 0.99 *±*0.15 *µ*m (vertical) (Fig. 1c(iii,iv)). The small deviation of *∼*0.1 *µ*m in the FWHM between the two systems lies within the pixel–sampling uncertainty and does not reflect a true optical performance difference, as the effective pixel sizes at the specimen plane are nearly identical, i.e., 0.323 *µ*m/px for the home-built microscope and 0.325 *µ*m/px for the commercial microscope. Fig. 1d summarizes the bead-by-bead analysis of the FWHM values as box plots. Example bead images and their corresponding horizontal and vertical line intensity profiles for both the home-built and commercial microscopes are shown (Supplementary Fig. 1). This bead-by-bead analysis captures the position-dependent measurement variability across the field of view. We exclude aggregated beads based on their visual appearance. The measured characteristics of individual beads vary with their location in the Si_3_N_4_ membrane and this effect is not accessible from averaged profiles alone.

In an ideal, aberration-free imaging system, the lateral point spread function is expected to be isotropic, yielding identical horizontal and vertical FWHM values for each bead, and consequently, a perfect linear correlation with *r* = 1.0. Deviations from this ideal case arise from residual optical imperfections such as weak astigmatism, alignment tolerances, or sampling effects introduced by the camera pixel size. Correlation analysis between the horizontal and vertical FWHM values of the beads shows a strong linearity in both microscopes of *r* = 0.80 for the home-built microscope and *r* = 0.76 for the commercial microscope (Fig. 1e). This high degree of correlation indicates that the lateral resolution in both systems is not dominated by axis-specific aberrations.

Finally, we validate the performance of the home-built microscope on a biological specimen, i.e., fluorescently labeled keratin bundles in fixed cells. Representative images acquired using the home-built (cyan) and commercial (magenta) microscopes include a line scan (A, dotted line) and a rectangular region of interest (ROI) (B, dotted box) for comparison (Fig. 1f (i,ii)). The normalized intensity profiles extracted along the line scan provide qualitative visualization of variation in fluorescence intensity along this line (Fig. 1f (iii)). However, since a line scan takes into account only limited number of pixels, the resulting profile is sensitive to shot noise, camera noise and precise placement of the line scan. For a further comparison of the two microscopes, we therefore extract normalized averaged intensity profiles from rectangular ROIs (Fig. 1f (iv)). The signal-to-noise ratio (SNR) and the contrast-to-noise ratio (CNR) are calculated from background-subtracted intensity profiles (B, dotted box region) using automatically detected signal peaks. The background level and noise are estimated from the lowest 20 % intensity values of the profile. For each detected peak, the mean signal intensity *µ*_signal_ is determined from a local region surrounding the peak (*±*3 pixels around the peak maximum). The SNR is then calculated as SNR = *µ*_signal_/*σ*_background_, where *σ*_background_ is the standard deviation of the background intensity. The CNR is calculated as CNR = (*µ*_signal_ *− µ*_background_)/*σ*_background_, where *µ*_background_ denotes the mean background intensity. The final SNR and CNR values are obtained by averaging over all detected peaks within the ROI. The home-built microscope exhibits an SNR of 21.2 compared to 15.5 for the commercial microscope, while the CNR is 11.15 for the home-built system and 9.42 for the commercial system.

Taken together, this bench marking shows that our compact home-built fluorescence microscope for integration into a synchrotron beamline achieves sub-micrometer optical resolution and an imaging performance comparable to state-of-the-art commercial research microscopes. At the same time, it satisfies the stringent mechanical, steric and operational constraints imposed by the synchrotron beamline environment. This enables precise correlative fluorescence and X-ray imaging with minimized time delay in-between.

### Design of the microfluidic chamber

Combined fast scanning SAXS and visible light fluorescence microscopy of fixed-hydrated and potentially living cells imposes stringent requirements on the sample environment. In particular, the cells must remain fully hydrated over extended measurement times, radiation-induced damage and gas bubble formation must be minimized, and the geometry must remain compatible with microscopy objectives for visible light fluorescence imaging. To enable correlative imaging of cells under fully hydrated, near-native conditions, we develop a dedicated microfluidic chamber compatible with both fast scanning SAXS and visible light fluorescence microscopy. Our chamber overcomes the above-mentioned limitations by combining a thin hydrated observation region with robust optical access and controlled perfusion.

Fig. 2a shows an overview of the experimental geometry. The microfluidic chamber is mounted on a scanning stage consisting of a hexapod for coarse positioning and a piezo stage for fine translations during both the microscopy and the SAXS measurements. During microscopy, the home-built epi-fluorescence microscope is moved along the y-axis and positioned in the X-ray path, thus aligning it with the microfluidic chamber (Fig. 2a(i)). During SAXS acquisition, the microscope is retracted along the y-axis, and the X-rays are focused on the cells in the microfluidic chamber (Fig. 2a(ii)). Importantly, this geometry avoids repositioning the chamber itself, enabling rapid alternation between the home-built microscope and scanning SAXS measurements and facilitating spatial registration between the two modalities. Further details of the integrated setup are provided in Experimental section .

**Figure 2:**
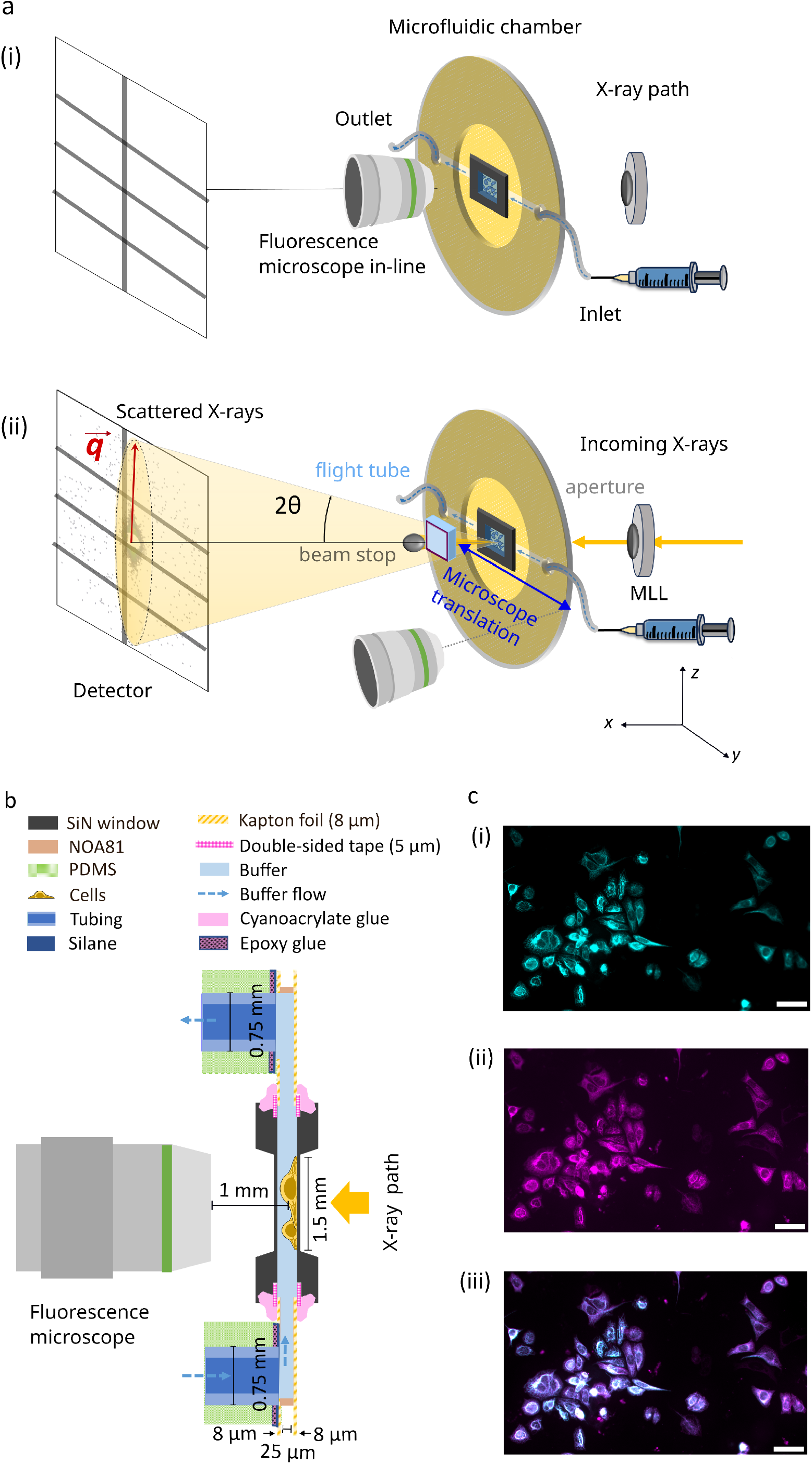
Design of the microfluidic chamber compatible with both fluorescence microscopy and X-ray measurements at the synchrotron beamline. (a) Schematic of the experimental setup (not to scale). (i) Microscopy mode: The home-built fluorescence microscope aligned with the microfluidic chamber for imaging. In this mode, no incoming X-ray beam is present. (ii) Fast scanning SAXS mode: The microscope is moved out of the X-ray path to enable fast scanning SAXS measurements. The incoming X-rays are focused by multilayer Laue lenses (MLLs), followed by an aperture placed downstream the MLLs, before illuminating the cells present in the microfluidic chamber. The primary beam is blocked by a beam stop and the scattered signal is collected by the detector. The red arrow illustrates the direction of the corresponding scattering vector 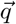 for a scattering angle (2*θ*). The biological cells are maintained in a thin hydration layer between two Si_3_N_4_ windows integrated in the microfluidic chamber, including an inlet (connected to a syringe) and an outlet for continuous buffer flow. (b) Side view schematic of the fully assembled microfluidic chamber and the geometry of the setup with the fluorescence microscope in-line while imaging. See legend for the components of the flow chamber and Supplementary Fig. 2 for a step-by-step explanation of the assembly. (c) Fluorescence images of fixed-hydrated keratin-rich SW-13 cells before (i, cyan) and after (ii, magenta) in situ immunostaining inside the assembled device. The overlay (iii) of the keratin structures before and after the in situ immunostaining shows good agreement. Scale bars: 50 *µ*m.

Fig. 2b shows a cross-sectional view of the fully assembled microfluidic chamber. Fabrication details are provided in Experimental section , with a step-by-step overview (Supplementary Fig. 2). The chamber consists of a Kapton–NOA81–Kapton device defining a 1.5 mm-wide microfluidic channel, with 25 *µ*m channel height. The central observation region is sealed from both sides by Si_3_N_4_ windows, thereby establishing an observation region that is compatible with both fast scanning SAXS and fluorescence microscopy. The channel dimensions are optimized to achieve sufficient volume flow rates at the observation site. This prevents sample drying and bubble formation caused by X-radiation. The channel dimensions also ensure the mechanical stability of the flow chamber and laminar flow conditions, as illustrated in the simulation (Supplementary Fig. 5). For additional mechanical stability, the device is mounted on a polydimethylsiloxane (PDMS) support containing a 3 cm diameter opening. This provides unobstructed access for a 20*×* , NA 0.7 objective. The enclosed water layer between the two Si_3_N_4_ membranes has a total thickness of approximately 60*±* 10 *µ*m.

This reduced water path length substantially decreases absorption, thereby improves the X-ray scattering contrast and enables exposure times as short as 2 ms. Prior to the Extremely Brilliant Source (EBS) upgrade at The European Synchrotron (ESRF), we performed measurements on fixed-hydrated SW-13 cells using a microfluidic chamber with a water layer of 200 *µ*m, lower photon flux, and longer exposure times [27]. A comparison of the corresponding dark field images is shown (Supplementary Fig. 6) and the experimental parameters before and after the EBS upgrade are summarized (Supplementary Table 2). Despite considerably shorter exposure times, we observe that the maximum dark-field contrast increases by nearly two orders of magnitude due to the higher photon flux. In addition, the thinner water layer enhances the scattering contrast, defined as (*I*_max_ *− I*_min_)/*I*_min_ by approximately a factor of two. Together, the reduced chamber thickness and higher photon flux enable rapid measurements with substantially enhanced scattering contrast without increasing the radiation dose. To maintain hydration and suppress X-ray-induced gas bubble formation, buffer is continuously perfused through the chamber during measurements. The experiments are typically conducted at a flow rate of 100 *µ*L/h. A summary of flow rates and corresponding velocities is provided (Supplementary Table 3). The chamber operates stably over a range of 20-200 *µ*L/h.

In addition to buffer perfusion, the microfluidic chamber enables controlled in situ delivery of reagents. To demonstrate this capability, we perform immunostaining of intracellular keratin networks inside the chamber. For this proof-of-principle experiment, fixed-hydrated SW-13 cells stably expressing fluorescent keratin are used, providing a reference for the quality of the immunostaining. Cells grown on Si_3_N_4_ membranes are integrated into the microfluidic chamber, followed by imaging the keratin structures (Fig. 2c(i,cyan)). This is followed by immunostaining including sequential perfusion steps as detailed in Experimental section. Following the staining, keratin structures are imaged again (Fig. 2c(ii, magenta)) and agree with the control image (Fig. 2c(iii, overlay)).

Taken together, the microfluidic chamber provides a well-controlled sample environment compatible with both visible light fluorescence microscopy and fast scanning SAXS. It combines stable perfusion, reduced water layer thickness, and suppression of radiation-induced gas formation. In addition, the chamber supports controlled in situ delivery of reagents, as demonstrated by immunostaining, providing a foundation for experiments on living cells related to drug addition or labeling. The platform is broadly applicable across cell types, staining strategies, and buffer conditions, and is compatible with both fixed and living cells.

### Multi-modal imaging of biological cells

The compact fluorescence microscope for beamline integration in combination with the dedicated microfluidic chamber enables sequential microscopy and fast scanning SAXS measurements of identical cellular regions under hydrated conditions. To demonstrate the versatility of this workflow, we apply the approach to two cell types with distinct cytoskeletal architectures: keratin bundles in SW-13 epithelial cells and force-generating actomyosin in hiPSC-derived cardiomyocytes.

SW-13 epithelial cells provide an ideal model system to investigate cytoskeletal organization due to their prominent and strongly bundled keratin K8/K18 intermediate filament network, which dominates the intracellular architecture. These cells stably express fluorescent keratin hybrids (HK8-CFP and HK18-YFP), enabling direct visualization of keratin organization without immunostaining. Fluorescence imaging reveals pronounced spatial heterogeneity within the keratin network (Fig. 3a). The inverted fluorescence image, shows wavy filament structures (black arrow) as well as straightened, highly ordered keratin bundles (black box) along with regions of dense keratin networks (red arrow). The corresponding X-ray dark-field image of the selected scan region is shown (Fig. 3b). The contrast between the cytoplasm (green) and nuclei (red) is well visible, indicating a good scattering contrast. This is achieved by integrating the scattering intensity over an optimized q-range, as described in Experimental section [20]. Notably, regions with dense keratin networks (Fig. 3a, red arrow) exhibit a higher scattering intensity (Fig. 3b) than those with lower keratin density.

**Figure 3:**
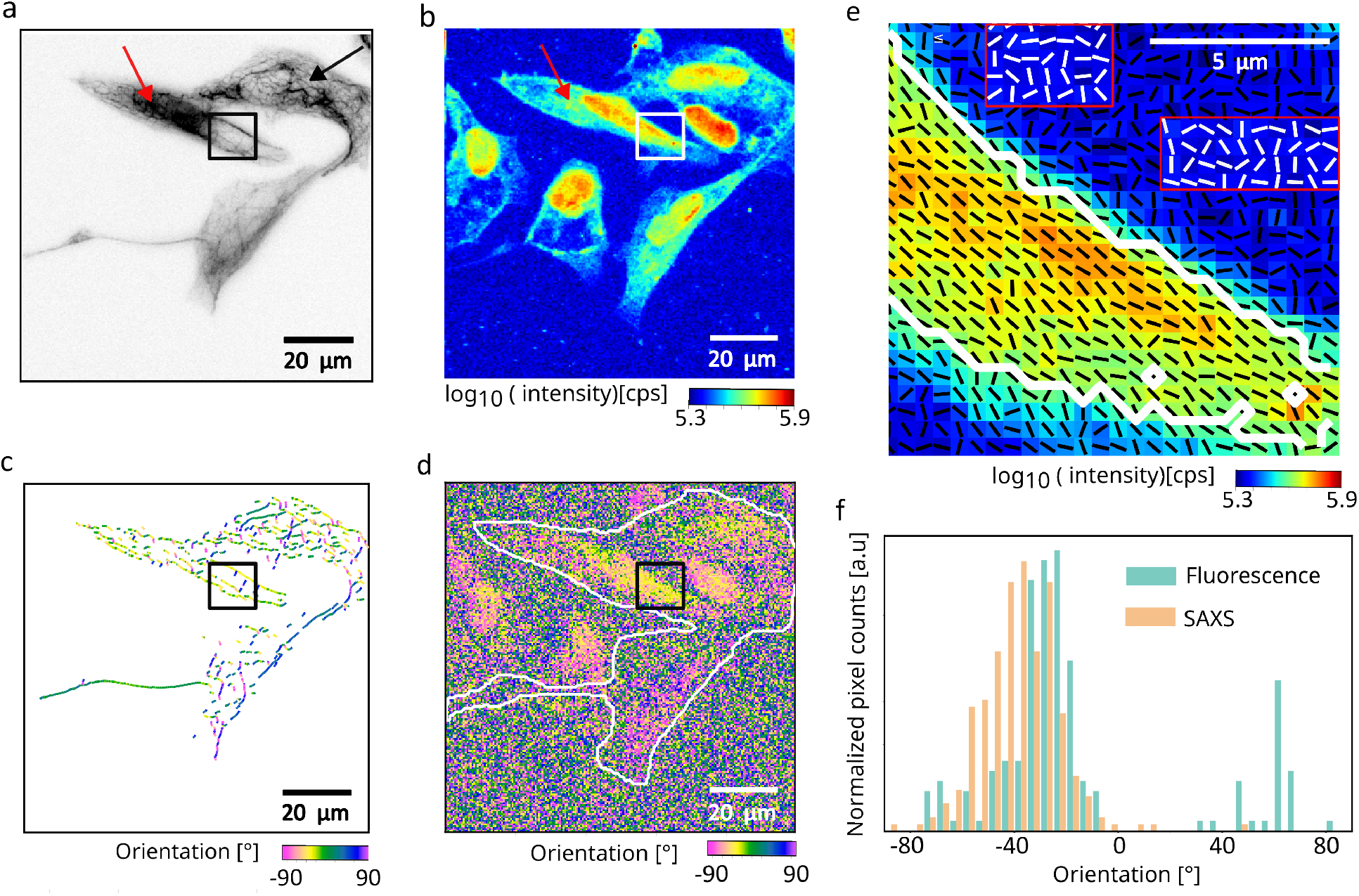
Correlative fluorescence and fast scanning SAXS analysis of keratin filament organization in SW-13 cells. (a) Inverted fluorescence image of keratin-rich SW-13 cells acquired using the home-built fluorescence microscope at the beamline. The black box highlights an area containing ordered keratin bundles, the black arrow points to wavy keratin structures, while the red arrow points to a dense keratin network. (b) X-ray dark-field image obtained from the scanning SAXS corresponding to the same region as shown in panel a. (c) Bundle networks are detected from the fluorescence image using SOAX and are color-coded by their orientation. (d) SAXS-derived orientation map; the white outlines indicate the keratin-expressing cells as shown in panel a. (e) Overlay of the X-ray dark-field image with black lines representing the SAXS-determined orientation of ordered structures as shown in the black box in panel a. Selected background regions are marked by red boxes, with white line segments indicating their corresponding orientations for enhanced visibility. Pixels within the white contour correspond to areas of high X-ray scattering intensity within the ordered structure. (f) Orientation histograms obtained from SAXS and fluorescence analysis for the same region within the white outline in panel d show consistent alignment trends between the two modalities.

In addition to the cells visible in the fluorescence image, the X-ray dark-field image reveals additional cells that do not express keratin and therefore lack fluorescence signal. This highlights the advantage of SAXS, where image contrast arises from electron density differences, independent of fluorescent labeling, enabling visualization of both labeled and unlabeled cellular components. Fig. 3c shows the keratin filament network detected from the fluorescence image using SOAX, with filaments color-coded according to their local orientation. The region containing highly ordered keratin bundles (black box in panel a), exhibits a consistent orientation, indicated by the yellow color corresponding to *∼* -30^*°*^.

Fig. 3d shows a scanning SAXS-derived orientation map, revealing local variations in filament alignment[20]. An orientation of 0^*°*^ corresponds to the horizontal direction (Fig. 3c,d). Regions containing ordered keratin bundles exhibit more uniform filament orientations, reflected by similar color values. Inter-estingly, both the orientation maps from fluorescence microscopy and from scanning SAXS show the same orientation distributions, see, e.g., the yellow color in the black box region. This result shows that both methods consistently detect oriented structures in the cells. Unlike fluorescence microscopy, SAXS is not specific to a particular labeled component and reports on the dominant orientational order arising from all anisotropic structures within the scanned region of the cell. A close inspection of the region highlighted by the black box (Fig. 3c, Fig. 3d), reveals agreement between the orientations obtained from fluorescence microscopy and SAXS, indicated by similar colors in both plots. This correspondence indicates that the keratin filament bundles considerably contribute to the SAXS-derived orientation.

For improved visualization, Fig. 3e shows an overlay of the X-ray dark-field image with black line segments for every scan pixel, representing the SAXS-derived local orientations. Here, the directions of the line segments indicate the directions of the cellular structures in real space and the lengths of the line segments correspond to the degree of anisotropy [13]. Some of the selected background regions are additionally marked by red contour boxes with white line segments to illustrate their random orientation patterns. The ordered keratin bundles identified in the fluorescence image (black box in Fig. 3a) correspond to coherent and well-defined SAXS orientations within the cell interior in the matching region of the dark-field image (white box in Fig. 3b). In contrast, regions outside the cell boundaries display random orientations. We further apply an X-ray intensity threshold to select pixels (white contour) corresponding to the cell regions with densely organized keratin structure, as observed in fluorescence image (black box in Fig. 3a). Only pixels with higher intensity values are included, since their SAXS-derived orientation values are considered more reliable due to improved signal-to-noise ratio. For quantitative comparison, the orientation histograms obtained from fluorescence microscopy and scanning SAXS are calculated within the white contour shown in panel e (Fig. 3f). The orientation values are determined on a pixel-wise basis from SOAX-processed fluorescence images and SAXS-derived orientation maps. To ensure spatial correspondence between the two modalities, the fluorescence image is first resampled onto the pixel grid of the X-ray dark-field image, followed by spatial registration, and the corresponding orientation values are considered for the histograms. The resulting orientation distributions from both imaging modalities show good agreement, with the peak of the distribution located around -30^*°*^. The strong agreement between the orientation distributions confirms that keratin filaments constitute the dominant ordered structures in SW-13 cells and demonstrates the utility of the multi-modal correlative imaging approach.

Ordered cytoskeletal assemblies are abundant across many cell types and provide distinct structural signatures that can be probed by complementary imaging modalities. While we first validated our multimodal approach on thick keratin bundles in epithelial cells, our approach is not limited to intermediate filaments. To demonstrate its broader applicability, we apply the multimodal imaging to the contractile actomyosin machinery in cardiomyocytes. Here, thin filaments made of actin and thick filaments made of myosin molecular motors slide past each other and thereby lead to contraction of the cells. These acto-myosin structures are anchored at the Z-discs, which are enriched in α-actinin. In our case, α-actinin is visualized via fusion with citrine fluorescent protein [28, 29]. Consequently, inverted gray scale fluorescence images of the cardiomyocytes clearly show the Z-discs as dark spots with periodic spacing (Fig. 4a). Note that due to the immature state of these iPSC cells, the Z-discs do not yet form well-defined transverse bands spanning the entire width of the myofibrils, as observed in the adult human cardiomyocytes. Instead, they appear as shorter and discontinuous structures. Fig. 4b shows the corresponding X-ray dark-field image of the selected scan region. A clear contrast between the cytoplasm (green) and nuclei (red) is well visible. Within the cytoplasm, regions exhibiting higher fluorescence intensity, i.e., regions with abundant actomyosin and Z-discs also show enhanced X-ray scattering intensity. Fig. 4c shows the orientation map derived from the fluorescence image shown in panel a by calculating the structure tensor and deriving the local orientation in each pixel by Eq. (3) in Experimental section. To validate the utility of the structure tensor for deriving an orientation map, we re-evaluate the orientation of the keratin bundles in the fluorescence images of the SW-13 cells (Fig. 3a, Supplementary Fig. 7). The orientation values derived from both SOAX and the structure tensor match well.

**Figure 4:**
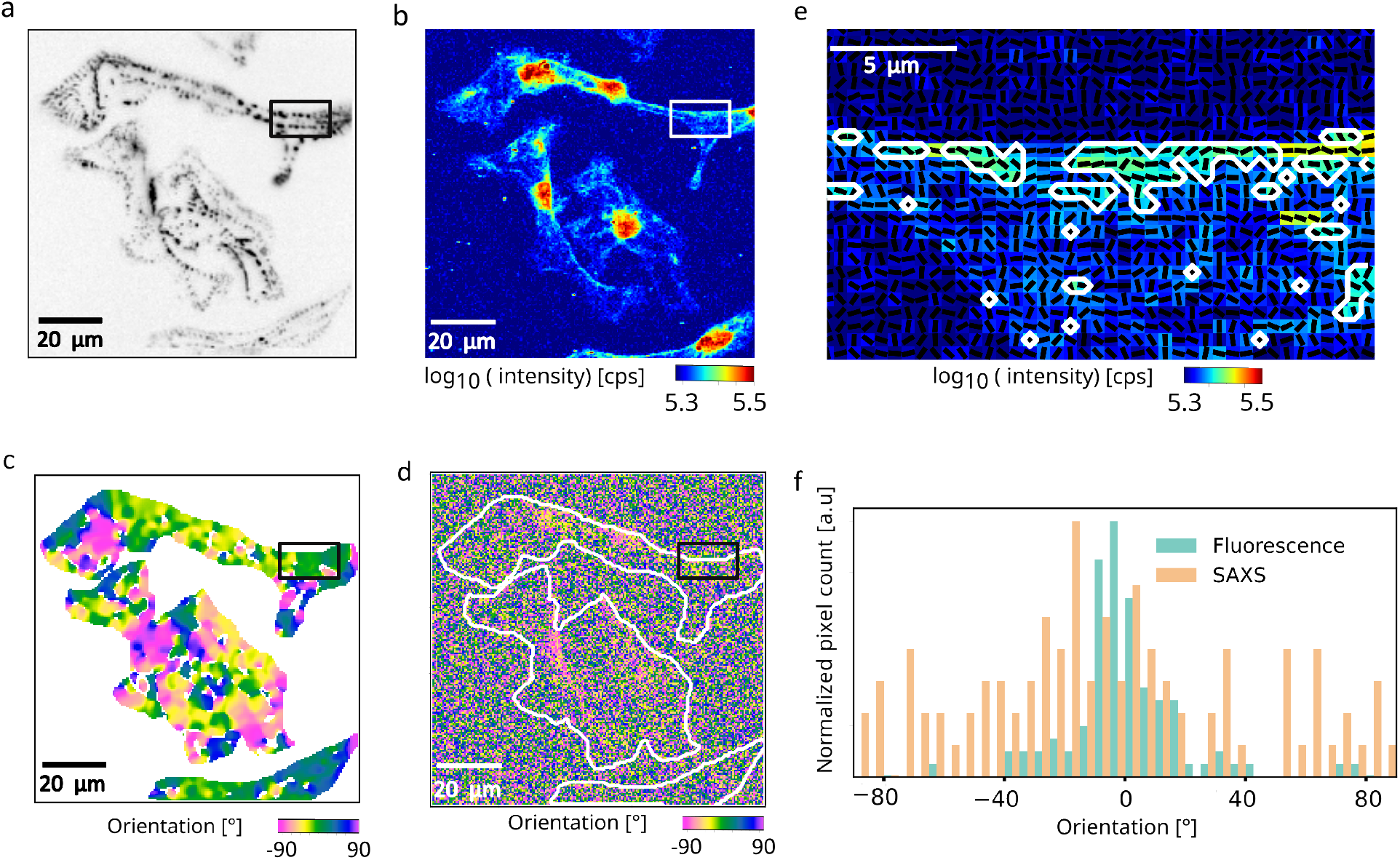
Correlative fluorescence and fast scanning SAXS analysis of sarcomeric organization in hiPSC-ACTN2-citrine-derived cardiomyocytes. (a) Inverted fluorescence image of α-actinin–labeled Z-discs with the periodic sarcomeric arrangement within the cardiomyocytes. The region within the black box highlights a representative area containing Z-discs. (b) The corresponding X-ray dark-field image reveals contrast differences between nucleus and the cytoplasm. (c) Orientation map derived from the fluorescence image of α-actinin–labeled Z-discs (d) SAXS-derived orientation map; the white outlines indicate the cells as shown in panel a. (e) Overlay of the X-ray dark-field image with black lines representing the SAXS-determined orientation of the region within the black box in panel a. Pixels within the white contour correspond to areas of high X-ray scattering intensity, where prominent Z-discs are observed in the fluorescence image. (f) Orientation histograms obtained from SAXS and fluorescence analysis for the same region within the white outline as shown in panel e.

The corresponding scanning SAXS-derived orientation map [20] of the same region is presented (Fig. 4d). The white outlines indicate the cell contours as obtained from the fluorescence image. We observe dominant orientation features in the regions of clearly resolved Z-discs as compared to the background region outside the white outline and cytoplasmic regions with low scattering intensity (see panel b). Notably, the SAXS-derived orientation reflects the alignment of anisotropic electron density modulations dominated by the highly ordered actomyosin structures. Identical color-coding is used for both modalities to facilitate direct comparison and, interestingly, the orientation maps from both the modalities show similar color distribution. To ensure meaningful quantitative comparison between the two modalities, we therefore select a region exhibiting a high degree of alignment from SAXS-derived orientation map (indicated by the box in panels a-d). In contrast to the keratin-rich SW-13 cells (Fig. 3), where dense filament bundles extend across large regions of the cytoplasm and give rise to strong X-ray scattering signals throughout the cell, the actomyosin organization in cardiomyocytes is spatially heterogeneous. As a result, the X-ray scattering intensity varies considerably across the cell, and regions with low scattering signal yield less reliable orientation estimates. We therefore restrict the quantitative comparison to regions exhibiting higher X-ray scattering intensity.

Fig. 4e shows an overlay of the X-ray dark-field image of the region in the box (panels a-d), with black line segments for every scan pixel representing the SAXS-determined orientation. The pixels within the white contours are selected based on their higher scattering intensity and correspond to regions containing Z-discs in the fluorescence image (panel b). These higher intensity pixels are included because they provide more reliable SAXS-derived orientation values.

For quantitative comparison, orientation histograms from fluorescence microscopy and SAXS are calculated within the white contour shown in panel e (Fig. 4f). The orientation values are obtained on a pixel-wise basis from the structure tensor applied to the fluorescence image and from the SAXS-derived orientation map after resampling of the fluorescence image into the pixel grid of the X-ray dark-field image, followed by spatial registration. The resulting orientation distributions from both the modalities show good agreement, with the peak of the distribution located around 0^*°*^. This is in line with what we expect for the immature Z-discs in hiPSCs as the structure tensor detects series of α-actinin clusters along the actomyosin structures. In mature cardiomyocytes one would expect more pronounced Z-discs elongate in perpendicular direction to the actomyosin, and, thus, an angular offset of 90^*°*^ between the distributions derived from the fluorescence images and the scanning SAXS data.

These results demonstrate that our multimodal imaging approach allows for a quantitative assessment of the degree of order in intracellular structures. Fluorescence microscopy provides molecular specificity, whereas scanning SAXS reveals the dominant structural alignment based on electron density contrast, thus highlighting the pronounced complementary of the two methods. The strong correlation between the independently derived orientation maps confirms that both methods consistently identify the same principal axis of filament organization, thereby validating the reliability of orientation analysis across imaging modalities.

## Discussion

Scanning SAXS is a powerful approach for probing structural organization in biological cells, revealing filament periodicities and bundle architecture with sub-micron precision [5–8, 13]. However, SAXS alone lacks molecular specificity and structural signals cannot be assigned to specific cellular components without complementary imaging. Correlative approaches have partially addressed this gap. Prior work combining cryo-fluorescence based techniques with cyro-soft X-ray tomography, whether using fiducial markers and separate imaging sessions or integrated on-line platforms [30, 31], has enabled the correlation between molecular identity and internal morphology of biological cells. Similarly, cyro-correlative fluorescence microscopy in combination with synchrotron X-ray fluorescence nanoimaging has provided access to elemental composition in frozen-hydrated cells [32, 33]. Correlative approaches specifically involving SAXS remain comparatively limited. The foundational work of Bernhardt et al. integrated STED fluorescence microscopy with scanning SAXS and X-ray holography, demonstrating the power of combining structural and molecular information, yet exclusively on freeze-dried samples, leaving the extension to hydrated and living biological systems, close to physiological condition, an open challenge [12, 25, 26].

The cytoskeleton is an ideal system for such investigation, as structure defines function in biological systems. Filamentous networks composed of intermediate filaments, actin filaments, and microtubules are routinely labeled for fluorescence microscopy, and their components exhibit a degree of distinct structural order that generates interpretable SAXS signals. Keratin-rich SW-13 cells and cardiomyocytes represent two systems with direct relevance to cell mechanics and force generation, making them a well suited test cases for a correlative platform.

Prior SAXS studies on SW-13 cells progressively refined understanding of keratin organization, from initial freeze-dried sample preparations establishing keratin bundle diameters on the order of tens of nm [13], through fixed-hydrated and living cell measurements in flow chambers with a 200-*µ*m water layer, to combined ptychography and scanning SAXS yielding quantitative bundle nanostructure [14]. However, all were limited by low photon flux, comparatively long acquisition time, and insufficient SAXS signal from hydrated samples and could not reliably extract orientation information. More recently, measurements on fixed-hydrated cells using a thin water layer (20 *µ*m) demonstrated fast scanning with milliseconds exposure times and enabled SAXS-derived orientation mapping through innovative analysis strategies [20]. However, molecular specificity through integrated fluorescence imaging remained missing.

For cardiomyocytes, correlative SAXS progressed from cryogenic to close-to-physiological conditions. On freeze-dried neonatal rat cardiomyocytes, home-laboratory fluorescence microscopy combined with scanning SAXS demonstrated orientation mapping [12], later extended to integrated STED microscopy and X-ray holography [25]. Moving to hydrated conditions, scanning SAXS on living iPS-cardiomyoyctes in static wet chambers produced X-ray darkfield images, while fixed-hydrated adult mouse cardiomyocytes yielded a detectable sarcomeric diffraction signature [34]. The two strongest equatorial reflections of actomyosin were later resolved, with orientation mapping achieved for fixed-hydrated sample preparations [18]. Later work probed sarcomeric diffraction in 3D iPS-cardiomyocytes tissue constructs under hydrated conditions [35], and meridional reflections, beyond equatorial peaks were detected in fixed-hydrated adult mouse cardiomyocytes [36]. Despite this progress, two gaps remained unaddressed: integrated fluorescence microscopy on the same sample in hydrated state was achieved so far and the orientation mapping in weakly ordered systems such as iPS-cardiomyocytes remained out of reach.

The present work closes both gaps. Our correlative platform integrates high-quality fluorescence microscopy with scanning SAXS in a configuration compatible with hydrated and living cells. The compact microscope hardware adapts to diverse beamline geometries without compromising image quality. The dedicated microfluidic chamber maintains an optimized water layer, thin enough for X-ray transmission, yet sufficiently thick to sustain cells and provide optical access in a physiologically relevant hydrated state. The same chamber enables chemical manipulation of samples during measurement, extending the scope of experiments that can be performed in a single session. Alongside these hardware developments, a dedicated data analysis framework enables quantitative correlation of the multimodal datasets.

The results validate the platform on two biologically relevant systems. In SW-13 cells, the correlative measurements confirm that keratin filaments constitute the dominant ordered structures contributing to the SAXS signal. The orientation distributions derived independently from fluorescence and from SAXS are in good agreement, establishing a clear link between the molecular identity and nanoscale structural organization. For fixed-hydrated iPS-cardiomyocytes, the results establish a direct correlation between fluorescence-resolved sarcomeric organization and SAXS-derived structural orientation. This is particularly notable given the structural immaturity characteristic of iPS-derived cardiomyocytes. Immature cardiomyocytes cultured on unpatterned substrates typically show comparatively isotropic structures, making orientation mapping challenging. Thus, this observed correlation supports the sensitivity of the platform and the robustness of the analysis strategy.

The platform opens several directions that were previously inaccessible. Time-resolved measurements on living cells under pharmacological or mechanical stimulation become feasible within the microfluidic chamber. Sequential fluorescence imaging with multiple filter sets can resolve several distinctly labeled components within the same cell, enabling richer multimodal datasets. Integration with additional X-ray modalities like holography, fluorescence or ptychography, is straightforward given the compact and adaptable instrument design. In a broader sense, the approach is not limited to the cytoskeletal systems demonstrated here, but it is applicable to any biological question where nanoscale structural organization and molecular identity must be addressed together under physiological conditions.

### Experimental

#### Cell culture and sample preparation

Human adrenal cortex carcinoma-derived SW-13 cells (ATCC CCL-105) [37] that stably express fluorescent keratin hybrids (HK8-CFP, HK18-YFP) [38, 39] are cultured in high-glucose Dulbecco’s Modified Eagle’s Medium (DMEM; 4.5 g/L; D6429, Merck, Darmstadt, Germany) with 10 % (v/v) fetal bovine serum (FBS; 10270-106, Gibco, Thermo Fischer Scientific, Waltham, MA, USA) and 1 % (v/v) penicillin–streptomycin (pen-strep, 100 U/mL penicillin and 0.1 g/L streptomycin; 15140122, Gibco), which we refer to as “SW-13 cell medium”. The cells are maintained at 37^*°*^C in a water-saturated atmosphere at 5 % CO_2_. After 1 week in culture, geneticin (L0015, Biowest, Nuaillé, France) is added (50 mg/mL) to the culture flask to suppress the cells which are not stably transfected. We refer to these cells as “SW-13 cells”.

In addition, a sarcomeric Z-band reporter hiPSC-line is used. This cell line has been developed by fusion of the C-terminus of α-actinin 2 gene with fluorescent protein citrine using CRISPR/Cas9 [28]. The differentiation of this hiPSC-line to the hiPSC-ACTN2 citrine derived cardiomyocytes is carried out following the protocol by Tiburcy et al. [40]. Cell culture flasks are coated with 0.8% (v/v) Matrigel (CLS354230-1EA, Merck) and are incubated at 37^*°*^C for 1 h prior to cell culture. The cells are cultured in Roswell Park Memorial Institute (RPMI) 1640 medium with GlutaMAX (61870010, Gibco), with 1% (v/v) pen-strep, 1 mmol/L sodium pyruvate and 2% B27 supplement (17504-044, Gibco), which we refer to as “CM medium”. Cultures are maintained at 37^*°*^C, 5% CO_2_, and the medium is changed every second day for at least two weeks before sample preparation. We refer to these cells as “cardiomyocytes”.

For sample preparation, silicon nitride (Si_3_N_4_) windows (frame size 5 *×*5 mm^2^, membrane size 1.5*×*1.5 mm^2^, thickness 1 *µ*m, Silson Ltd, Warwickshire, UK) are plasma-treated (50 W, 30 s; Zepto, Diener electronic GmbH & Co. KG, Ebhausen, Germany) and UV-sterilized. For plating the SW-13 cells, the windows are coated with 5 % (v/v) fibronectin (F1141, Merck) and incubated at 37^*°*^C for 1 h. The cells cultured in the flasks are briefly washed with Dulbecco’s phosphate buffered saline (DPBS; D8537, Merck) and detached using 0.25% (v/v) Trypsin - 0.02% (w/v) EDTA (P10-020500, PAN-Biotech GmbH, Aidenbach, Germany). The cells are seeded at a density of 1.5 *×*10^5^ cells/mL on the treated Si_3_N_4_ windows. After 24 h, the cells grown on the Si_3_N_4_ windows are fixed with 4 % methanol-free formaldehyde (28906, Thermo Fisher Scientific). For plating the cardiomyocytes, the Si_3_N_4_ windows are coated with 0.8% (v/v) Matrigel and incubated at 37^*°*^C for 1 h. The cells cultured in the flask are briefly washed with DPBS, detached using Accutase (A11105-01, Gibco) with 0.025 % Trypsin (15090-046, Gibco) and 20 *µ*g/mL DNase I (260913, Calbiochem-Merck), and seeded at 1.5 *×*10^5^ cells/mL on the treated Si_3_N_4_ windows. The cultures are maintained for 18 h in CM medium supplemented with 5 *×*10^*−*6^ mol/L Y-27632-2HCl (orb154626, Biorbyt, Durham, NC, USA), followed by only CM medium up to 5 days before fixation with 4% methanol-free formaldehyde. The fixed cells on Si_3_N_4_ windows are maintained in DPBS to ensure continuous hydration.

#### Fabrication of the microfluidic chamber

A silicon master for the chamber fabrication with a flow channel and supporting structures, made from SU-8, is fabricated by standard photolithography methods (Supplementary Fig. 3a). Briefly, a 2-inch silicon wafer (MicroChemicals, Ulm, Germany) is spin coated with SU8 negative photoresist (SU-8 3025; MicroChem, Newton, MA, USA) to achieve a height of 25 *µ*m. The resist is then exposed to UV light through a photomask (Selba, Versoix, Switzerland) containing the dedicated structure design and is developed (Supplementary Fig. 2a). The silicon master is coated with fluorosilane (1H,1H,2H,2H-perfluorooctyltriethoxysilane, 97%; AB104055, abcr GmbH, Karlsruhe, Germany) by overnight vapor deposition. A PDMS replica for device fabrication is made by mixing polydimethylsiloxane (PDMS; Sylgard 184, Biesterfeld Spezialchemie GmbH, Hamburg, Germany) base and curing agent at a standard ratio of 10:1 (w/w) and curing the mixture on the silicon master (Supplementary Fig. 2b). Consequently, a single channel with a width of 1500 *µ*m and a height of 25 *µ*m surrounded by repeating cylinder structures of 700 *µ*m diameter with the center-center distance of 1000 *µ*m is obtained in the PDMS replica. These cylindrical structures locally alter the capillary pressure, guiding fluid flow and enabling precise control of capillary filling of the structure by Norland Optical Adhesive 81 (NOA 81, Norland Products, Jamesburg, NJ, USA) in the next step. To do so, the PDMS replica is placed on a Kapton foil (8 *µ*m thickness, SPEX SamplePrep, Metuchen, NJ, USA) on a flat aluminum block to serve as the bottom substrate. 80 *µ*L NOA 81 are pipetted around the PDMS replica (Supplementary Fig. 2c). After the capillary filling, the NOA 81 is cured under a UV lamp (365 nm, 2 x 8 W tubes; Herolab GmbG Herolab GmbH Laborgeräte, Wiesloch, Germany) for 2 minutes, and the PDMS replica is removed (Supplementary Fig. 2d). A second Kapton foil is mounted onto a microscope slide (75*×* 50 mm^2^, thickness 0.96 to 1.06 mm, Corning Incorporated, NY, USA) using adhesive tape at the edges to ensure flatness. The Kapton surface is plasma activated for 30 s at 50 W (Zepto plasma cleaner, Diener electronic GmbH & Co. KG, Ebhausen, Germany), and adhered to the NOA 81–Kapton assembly, to form the Kapton–NOA 81–Kapton device (Supplementary Fig. 2e). The device is baked at 65^*°*^C for 16 hours under light weight (*∼* 30 g) to enhance the bonding strength and ensure proper contact (Supplementary Fig. 2f).

To provide mechanical support of the device, we bond it to a ring–shaped piece of PDMS (inner diameter 3 cm, outer diameter 5.08 cm, thickness *∼*1 cm). The PDMS support is attached to the device using a thiolepoxy-click reaction [41]. To do so, the PDMS is plasma activated for 30 s at 50 W, and immediately placed under a desiccator with 40 *µ*L of mercaptosilane (3-Mercaptopropyl trimethoxysilane; 175617, Sigma-Aldrich, Merck) on a glass cover slip for vapor deposition for at least 2 h. Following the coating, a two-component epoxy glue (UHU 45705, Bühl, Germany) is applied to the device, and the coated PDMS support is adhered (Supplementary Fig. 2g). In order to generate a stronger bond, the assembled chamber is compressed under low weight (*∼* 30 g) for at least 3 h. Following the fabrication, a 3.5 mm hole is punched at the center of the flow channel using a biopsy puncher (Plano, Wetzlar, Germany), and one empty Si_3_N_4_ window is attached using thin double-sided adhesive tape (5 *µ*m; Nitto, Osaka, Japan). Holes are also punched at the inlet and outlet positions of the microfluidic device with a 0.75 mm diameter puncher, followed by insertion of tubing (inner diameter: 0.38 mm, outer diameter: 1.09 mm, polyethylene; Becton, Dickinson and Company, NJ, USA) pre-filled with Dulbecco’s phosphate buffered saline (DPBS, Merck)(Supplementary Fig. 2h). The tubing is preconnected to a gas tight syringe (2.5 mL, PTFE Luer Lock; Hamilton, Reno, NV, USA) to avoid enclosing of air bubbles. The channel is checked for free fluid flow without clogging using a microfluidic pump (Landgraf Laborsysteme HLL GmbH, Langenhagen, Germany) at a flow rate of 100 *µ*L/hr. A Si_3_N_4_ window with cells grown on it is rinsed with ultrapure water to remove remaining salt or dust particles. The frame area around the window is gently blotted, followed by careful alignment of the frame and membrane area with the respective areas of the empty Si_3_N_4_ window, using double-sided adhesive tape. Additionally, a small amount of cyanoacrylate adhesive (UHU 478747-62, Bühl, Germany) is added around the frame edges of both Si_3_N_4_ windows for sealing, resulting in the fully assembled microfluidic chamber (Supplementary Fig. 2i). During the bonding process, a constant flow rate of 20 *µ*L/hr is maintained to prevent dehydration of the cells. Once the glue is cured completely, the flow rate is increased to 100 *µ*L/hr and the parts of the chamber holder are assembled (Supplementary Fig. 3c).

#### Immunostaining within the microfluidic chamber

Following the microfluidic chamber assembly, immunostaining is initiated by supplying staining solutions via the inlet. Since the chamber is designed with a single inlet and outlet, the staining solutions are supplied sequentially by exchanging the gastight syringe connected to the inlet. To minimize the introduction of air bubbles during syringe exchange, only the syringe is replaced while keeping the needle–tubing assembly in place. We begin with perfusion of 1% Triton X-100 (3051.2, Carl Roth, Karlsruhe, Germany) in DPBS at a flow rate of 50 *µ*L/hr to permeabilize the cell membranes. The Triton solution reaches the cell window after *∼*25 minutes. The inlet is then immediately switched to DPBS and rinsed at 100 *µ*L/h for 30 minutes. The DPBS reaches the cell window after *∼*12.5 minutes, providing a net Triton incubation time of *∼*12.5 minutes. To block non-specific binding, 3% bovine serum albumin (130-091-376, MACS BSA Stock Solution; Miltenyi Biotec) in DPBS is perfused at 100 *µ*L/h for 30 minutes. The flow is then stopped, and the sample is incubated under static conditions for 45 minutes. Following incubation with the BSA solution, a primary antibody against K8 keratin, Troma I (0.5 *µ*g/mL in 3% BSA; AB531826, DSHB, Iowa, USA), is introduced into the chamber at a flow rate of 50 *µ*L/h for 40 minutes. The flow is then stopped, and the sample is incubated for 1.5 hours under static conditions. The chamber is subsequently rinsed with DPBS at 100 *µ*L/h for 30 minutes. Secondary antibody staining is performed using 1 *µ*g/mL goat anti-rat IgG conjugated with Alexa Fluor Plus 647 (A32733TR, Thermo Fisher Scientific), which is perfused at 50 *µ*L/h for 30 minutes. The flow is again stopped, and the sample is incubated for an additional hour. Finally, the sample is rinsed with DBPS at a flow rate of 100 *µ*L/h for 30 minutes. All imaging before and after immunostaining is performed on an inverted microscope (IX83, Olympus/Evident) using an IX3-FGFPXL filter set (excitation: 460–480 nm; emission: 495–540 nm; dichroic: 490 nm; Olympus/Evident), an ET-Cy5 filter set (excitation: 590–650 nm; emission: 662–738 nm; dichroic: 660 nm; Olympus/Evident) and a 20*×* objective (NA 0.7, UCPLFLN20X, Olympus/Evident).

#### Integrated fast scanning SAXS and fluorescence microscopy

The combined fast scanning SAXS and visible light fluorescence microscopy experiments are performed at the nanobranch (experimental hutch 3, EH3) of the ID13 beamline at the European Synchrotron (ESRF, Grenoble, France). The workflow comprises the microfluidic chamber assembly, identification of regions of interest (ROIs) based on fluorescence microscopy, and targeted scanning SAXS measurements of the same regions. Fluorescence microscopy is used to select cells exhibiting ordered cytoskeletal structures. The keratin structures in SW-13 cells and the Z-discs in cardiomyocytes are imaged with a YFP filter-set and 100 ms exposure time (Supplementary Table 1). The motor positions of each ROI are stored to ensure repositioning during scanning SAXS measurements.

Prior to scanning SAXS measurements, the microscope is retracted along the y-axis by *∼*125 cm to clear the beam path. The undulator beam is monochromatized by a channel-cut Si (111) monochromator to a photon energy of 15.0 keV. The beam is prefocused by beryllium parabolic refractive lenses (Be-CRLs). The X-rays are focused by a multilayer Laue lens (MLL; focal length 90 mm, from ESRF long term proposal collaboration MI-1463)) and spatially cleaned by a square shaped 40 *×*40 *µ*m^2^ PtIr aperture (order-sorting aperture, OSA), resulting a beam spot size of about 250*×* 250 nm^2^ (horizontal × vertical) at the sample position inside the microfluidic chamber. A helium-filled flight tube is placed between the microfluidic chamber and the SAXS beamstop (made of lead, *∼*250 *µ*m diameter), leaving only a few millimeters of air path on both sides of the flight tube. This configuration helps minimizing the background contribution from air scattering. The X-ray scattering patterns are recorded using an Eiger X 4M detector (Dectris, Baden, Switzerland) with 2070 *×*2167 pixels^2^ (4 megapixels) and a pixel size of 75 *×*75 *µ*m^2^ located at a sample-to-detector distance of about 1.9 m. This distance is calibrated using AgBeh powder placed in a empty chamber. In fast scanning SAXS mode, data are acquired using continuous line scans with constant stage motion. A 100 *×*100 *µ*m^2^ scan area with 0.5 *µ*m *×*0.5 *µ*m step size (40,000 scattering patterns) requires *∼* 4 min at 5 ms exposure per position (used for the cardiomyocytes) or *∼*3 min at 2 ms exposure per position (used for the SW-13 cells) At the end of each scan line, the recording is paused while the piezo stage moves to the start of the next line, while the sample is still being exposed.

## Data Analysis

### SAXS data

We follow the analysis method described by Yu et al. [20]. The X-ray dark-field image is obtained by integrating the total scattering intensity within an optimized q-range of 0.19–0.39 nm^*−*1^ for each scattering pattern. This approach effectively denoises the data and enhances contrast in the dark-field image. From the collected scattering patterns, the local orientation is computed by evaluating the circular mean of the azimuthal intensity distribution after background subtraction and masking. This analysis yields spatially resolved maps of the predominant scattering orientation, representing the local structural organization within the sample.

### Spatial image registration

The fluorescence images are rescaled by interpolation to match the pixel size of the X-ray dark-field images to ensure a common spatial sampling grid. Image registration is performed on the rescaled fluorescence image and the X-ray dark-field image based on cross-correlation in Fourier space [42]. The displacement (Δ*y*, Δ*x*) is computed with subpixel precision using an upsampling factor of 20. The detected translation is applied using the Fourier shift theorem. In this framework, a spatial shift in real space corresponds to a phase modulation in Fourier space, I^*′*^(**q**) = I(**q**)*e*^*−i***q**· Δ**r**^ where I(**q**) denotes the Fourier transform of the image and Δ**r** denotes the displacement. The registered image is obtained by inverse Fourier transformation.

### Filament-tracing based orientation mapping

The orientation of the keratin bundles in SW-13 cells as observed in the fluorescence images is quantified by filament detection followed by azimuthal angle analysis. The filaments are traced using the software package SOAX (version 3.7.0) [43]. We set the parameter “init-z” to false, set the ridge-threshold to 0.008 and the minimum filament length to 10 pixels. All other parameters are set to the default values. The traced filament coordinates are exported and further analyzed using a custom Python script to determine local filament orientations. For each filament detected by SOAX, the azimuthal angle is calculated from the local tangent direction between consecutive filament points. Specifically, for each segment defined by two neighboring coordinates (x_1_,y_1_) and (x_2_,y_2_) the in-plane angle is computed as

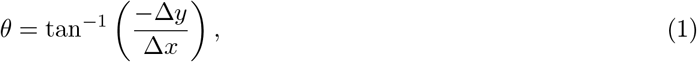

where Δ*x* = *x*_2_ *− x*_1_, and Δ*y* = *y*_2_ *− y*_1_. This angle is then mapped onto the symmetric range [-90^*°*^, +90^*°*^] to represent orientations (i.e., directions without polarity). Hence, the azimuthal angle corresponds to the local in-plane orientation of each filament segment with respect to the horizontal axis (*x*-axis) of the image, i.e., the horizontal x-axis is considered as 0^*°*^, and the vertical *y*-axis is considered to be +90^*°*^.

### Structure-tensor based orientation mapping

We apply a structure tensor based approach [44] to evaluate the orientation of keratin filaments in SW-13 cells and labeled Z-discs in cardiomyocytes. Unlike filament tracing methods, which require continuous filamentous structures, the structure tensor estimates local orientations directly from image intensity gradients and therefore does not rely on explicit segmentation. This makes it particularly suitable for structures like Z-discs. For an image intensity distribution *I(x, y)*, the spatial intensity gradients 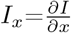 and 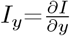 are computed after Gaussian presmoothing with a gradient scale *σ*. The local structure tensor is defined as

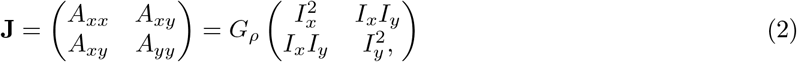

where, *G*_*ρ*_ denotes Gaussian averaging with the integration scale *ρ*. The structure tensor components, (*A*_*xx*_, *A*_*xy*_, *A*_*yy*_) thus represent locally averaged second-order intensity gradients. The local orientation angle is computed as

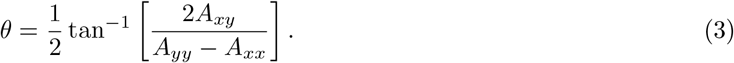

*θ* corresponds to the direction of the dominant eigenvector of the structure tensor and yields the local orientation of image features. For our analysis, the gradient scale is set to *σ* = 1.0 pixels and the integration scale is set to *ρ* = 2.0 pixels. The gradient scale determines the spatial extent of structures contributing to the derivative, whereas the integration scale controls spatial averaging of the tensor components, therefore improving robustness against noise. To restrict the analysis to regions exhibiting a dominant orientation, we compute the local anisotropy from the eigenvalues (*λ*_1_ *≥ λ*_2_) of the structure tensor **J** as

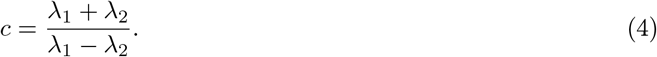

This value ranges from 0 (isotropic, no preferred direction) to 1 (highly oriented). Pixels with anisotropy < 0.2 are excluded to guarantee reliable orientation estimation. To demonstrate the utility of the approach, we apply this procedure to synthetic data showing stripe patterns with different orientations (horizontal, vertical and diagonal) (Supplementary Fig. 8).

### Orientation histograms

Pixel-wise orientation values are extracted from the spatially registered fluorescence images and the SAXS-derived orientation maps. For quantitative comparison, both orientation distributions are normalized such that the integral equals 1. The normalized data are rebinned over the angular range [-90^*°*^, +90^*°*^] into 36 bins (5^*°*^ width) by weighted integration within each interval, followed by renormalization. The bin centers, defined as the midpoints of the angular intervals, are used to represent the mean orientation in the comparison plots.

## Acknowledgements

We are grateful for the long term proposal beam time (SC-5436) granted by the ESRF (The European Synchrotron, Grenoble, France), and the excellent support of the ID 13 staff and the PSCM (Partnership for Soft Condensed Matter), specifically Pierre Lloria, and Peter van der Linden. We thank Peter Gawlitza and Sven Niese for providing MLL-lens prototypes used for these experiments and Wonhee Lee for helpful suggestions on the microfluidic chamber design. We further thank Claudia Geisler for fruitful discussions regarding microscopy. This work was financially supported by the German Federal Ministry of Research, Technology and Space (BMFTR) under grants No. 05K22MG3 and 05K25MGA (to S.K.) and the German Research Foundation (DFG): project-ID 449750155 - RTG 2756, projects A2, A7 (to S.K.) and under Germany’s Excellence Strategy - EXC 2067, ‘Multiscale Bioimaging: from Molecular Machines to Networks of Excitable Cells’ (MBExC, grant No. EXC 2067/1-390729940, to S.K. and W.H.Z.) and in the framework of DAPHNE4NFDI (grant No. 460248799), to S.K.. W.H.Z. is additionally supported by the DZHK (German Center for Cardiovascular Research), the German Federal Ministry for Science and Education (BMBF FKZ 161L0250A), and the Foundation Leducq (20CVD04).

## References

(1) Hémonnot, C. Y. J.; Köster, S. ACS Nano 2017, 11, 8542–8559.

(2) Albers, J.; Svetlove, A.; Duke, E. J. Cell Sci. 2024, 137, jcs261953.

(3) Fantoni, S.; Brombal, L.; Cardarelli, P.; Baruffaldi, F. Proc. R. Soc. A: Math. Phys. Eng. Sci. 2025, 481, 20250500.

(4) Priebe, M.; Bernhardt, M.; Blum, C.; Tarantola, M.; Bodenschatz, E.; Salditt, T. Biophys. J. 2014, 107, 2662–2673.

(5) Lichtenegger, H.; Reiterer, A.; Stanzl-Tschegg, S.; Fratzl, P. J. Struct. Biol. 1999, 128, 257–269.

(6) Fratzl, P.; Jakob, H. F.; Rinnerthaler, S.; Roschger, P.; Klaushofer, K. J. Appl. Crystallogr. 1997, 30, 765–769.

(7) Rinnerthaler, S.; Roschger, P.; Jakob, H. F.; Nader, A.; Klaushofer, K.; Fratzl, P. Calcif. Tissue Int. 1999, 64, 422–429.

(8) Gourrier, A.; Li, C.; Siegel, S.; Paris, O.; Roschger, P.; Klaushofer, K.; Fratzl, P. J. Appl. Crystallogr. 2010, 43, 1385–1392.

(9) Tesch, W.; Eidelman, N.; Roschger, P.; Goldenberg, F.; Klaushofer, K.; Fratzl, P. Calcif. Tissue Int. 2001, 69, 147–157.

(10) Gaiser, S.; Deyhle, H.; Bunk, O.; White, S. N.; Müller, B. Biointerphases 2012, 7.

(11) Fratzl, P.; Klaushofer, K.; Gupta, H. S.; Roschger, P.; Zizak, I.; Fratzl-Zelman, N.; Nader, A. Calcif. Tissue Int. 2003, 72, 567–576.

(12) Bernhardt, M.; Nicolas, J.-D.; Eckermann, M.; Eltzner, B.; Rehfeldt, F.; Salditt, T. New J. Phys. 2017, 19, 013012.

(13) Weinhausen, B.; Nolting, J.-F.; Olendrowitz, C.; Langfahl-Klabes, J.; Reynolds, M.; Salditt, T.; Köster, S. New J. Phys. 2012, 14, 085013.

(14) Hémonnot, C. Y. J.; Reinhardt, J.; Saldanha, O.; Patommel, J.; Graceffa, R.; Weinhausen, B.; Burghammer, M.; Schroer, C. G.; Köster, S. ACS Nano 2016, 10, 3553–3561.

(15) Piazza, V.; Weinhausen, B.; Diaz, A.; Dammann, C.; Maurer, C.; Reynolds, M.; Burghammer, M.; Köster, S. ACS Nano 2014, 8, 12228–12237.

(16) Hémonnot, C. Y. J.; Ranke, C.; Saldanha, O.; Graceffa, R.; Hagemann, J.; Köster, S. ACS Nano 2016, 10, 10661–10670.

(17) Weinhausen, B.; Saldanha, O.; Wilke, R. N.; Dammann, C.; Priebe, M.; Burghammer, M.; Sprung, M.; Köster, S. Phys. Rev. Lett. 2014, 112, 088102.

(18) Reichardt, M.; Neuhaus, C.; Nicolas, J.-D.; Bernhardt, M.; Toischer, K.; Salditt, T. Biophys. J. 2020, 119, 1309–1323.

(19) Weinhausen, B.; Köster, S. Lab Chip 2013, 13, 212–215.

(20) Yu, B.; Sinha, M.; Da Silva, R. M.; Rölleke, U.; Burghammer, M.; Köster, S. J. Synchrotron Radiat. 2026, 33.

(21) Köster, S.; Pfohl, T. Mod. Phys. Lett. B. 2012, 26, 1230018.

(22) Ghazal, A.; Lafleur, J. P.; Mortensen, K.; Kutter, J. P.; Arleth, L.; Jensen, G. V. Lab Chip 2016, 16, 4263–4295.

(23) Cassini, C.; Wittmeier, A.; Brehm, G.; Denz, M.; Burghammer, M.; Köster, S. J. Synchrotron Radiat. 2020, 27, 1059–1068.

(24) Xia, Y.; Whitesides, G. M. Angew. Chem. Int. Ed. 1998, 37, 550–575.

(25) Bernhardt, M.; Nicolas, J.-D.; Osterhoff, M.; Mittelstädt, H.; Reuss, M.; Harke, B.; Wittmeier, A.; Sprung, M.; Köster, S.; Salditt, T. Nat. Commun. 2018, 9, 3641.

(26) Bernhardt, M.; Nicolas, J.-D.; Osterhoff, M.; Mittelstädt, H.; Reuss, M.; Harke, B.; Wittmeier, A.; Sprung, M.; Köster, S.; Salditt, T. J. Synchrotron Radiat. 2019, 26, 1144–1151.

(27) Weinhausen, B. Scanning X-Ray Nano-Diffraction on Eukaryotic Cells: From Freeze-Dried to Living Cells, PhD thesis, Georg-August-Universität Göttingen, 2013.

(28) Haertter, D.; Hauke, L.; Driehorst, T.; Nishi, K.; Zimmermann, W.-H.; Schmidt, C. F. 2026, DOI: https://www.biorxiv.org/content/10.1101/2024.05.28.596183v2.

(29) Härtter, D.; Hauke, L.; Driehorst, T.; Long, Y.; Bao, G.; Primeßnig, A.; Berečić, B.; Cyganek, L.; Tiburcy, M.; Schmidt, C. F.; Zimmermann, W.-H. 2025, DOI: http://www.biorxiv.org/content/early/2025/05/04/2025.04.29.650605.

(30) Smith, E. A.; Cinquin, B. P.; Do, M.; McDermott, G.; Gros, M. A. L.; Larabell, C. A. Ultramicroscopy 2014, 143, 33–40.

(31) Zhang, C.; Guan, Y.; Tao, X.; Tian, L.; Chen, L.; Xiong, Y.; Liu, G.; Wu, Z.; Tian, Y. Opt. Express 2024, 32, 27508–27518.

(32) Karpov, D.; Cuau, L.; Shishkov, R.; Gramaccioni, C.; Dallerba, E.; Schwehr, B. J.; Ellison, G.; Hackett, M. J.; Plush, S. E.; Massi, M.; Lerouge, F.; Cloetens, P.; Bohic, S. ACS Nano 2026, 20, 7401–7413.

(33) Ortega, R.; Fernández-Monreal, M.; Pied, N.; Roudeau, S.; Cloetens, P.; Carmona, A. Chem. Biomed. Imaging 2024, 2, 744–754.

(34) Nicolas, J.-D.; Bernhardt, M.; Schlick, S. F.; Tiburcy, M.; Zimmermann, W.-H.; Khan, A.; Markus, A.; Alves, F.; Toischer, K.; Salditt, T. Prog. Biophys. Mol. Biol 2019, 144, 151–165.

(35) Javor, J.; Ewoldt, J. K.; Cloonan, P. E.; Chopra, A.; Luu, R. J.; Freychet, G.; Zhernenkov, M.; Ludwig, K.; Seidman, J. G.; Seidman, C. E.; Chen, C. S.; Bishop, D. J. Microsyst. Nanoeng. 2021, 7, 10.

(36) Bruns, H.; Czajka, T. S.; Sztucki, M.; Brandenburg, S.; Salditt, T. Biophys. J. 2024, 123, 3024–3037.

(37) Leibovitz, A.; McCombs, W. M.; Johnston, D.; McCoy, C. E.; Stinson, J. C. J. Natl. Cancer Inst. 1973, 51, 691–697.

(38) Strnad, P.; Windoffer, R.; Leube, R. E. J. Cell Sci. 2002, 115, 4133–4148.

(39) Windoffer, R.; Woll, S.; Strnad, P.; Leube, R. E. Mol. Biol. Cell 2004, 15, 2436–2448.

(40) Tiburcy, M. et al. Circulation 2017, 135, 1832–1847.

(41) Hoang, M. V.; Chung, H.-J.; Elias, A. L. J. Micromech. Microeng. 2016, 26, 105019.

(42) Guizar-Sicairos, M.; Thurman, S. T.; Fienup, J. R. Opt. Lett. 2008, 33, 156–158.

(43) Xu, T.; Vavylonis, D.; Tsai, F.-C.; Koenderink, G. H.; Nie, W.; Yusuf, E.; I-Ju Lee; Wu, J.-Q.; Huang, X. Sci. Rep. 2015, 5, 9081.

(44) Rezakhaniha, R.; Agianniotis, A.; Schrauwen, J. T. C.; Griffa, A.; Sage, D.; Bouten, C. V. C.; van de Vosse, F. N.; Unser, M.; Stergiopulos, N. Biomech. Model. Mechanobiol. 2012, 11, 461–473.

