## Supplementary Information for "Correlative X-ray imaging and fluorescence microscopy"

Table 1: List of components used to construct the home-built fluorescence microscope set-up, sorted along the light path

| **Component** | **Model** | **Company** | **Specifications** |
|---|---|---|---|
| LED light source | X-Cite Xylis II, XT730L | Excelitas Technologies Pittsburgh, PA, USA | Emission spectrum: 360 nm – 770 nm |
| Light guide | – | Excelitas Technologies | Diameter: 3 mm;<br>Length: 1.5 m |
| Aspheric collimating lens | ACL2520U-A | Thorlabs, Newton, NJ, USA | Focal length $f = 30$ mm;<br>Numerical aperture NA = 0.6 |
| Kinematic cage system | DFM1T1 | Thorlabs | 30 mm cube for filter set |
| 20× objective | UCPLFLN20X | Evident, Hamburg, Germany | NA = 0.7;<br>Working distance WD = 0.8 – 1.8 mm;<br>Adjustable cover glass correction 0 – 1.6 mm |
| GFP filter set | L70-450GF | AHF, Tübingen, Germany | Excitation filter: 460/50 nm;<br>Dichroic mirror: 495 nm;<br>Emission filter: 525/50 nm |
| YFP filter set | F36-529 | AHF | Excitation filter: 504/24 nm;<br>Dichroic mirror: 521 nm;<br>Emission filter: 539/27 nm |
| Tube lens | TTL 180A | Thorlabs | $f = 180$ mm;<br>WD = 133.5 mm;<br>anti reflection coating: 350 - 700 |
| Camera | pco.pixelfly 1.4 M-USB | Excelitas Technologies | 1392 × 1040 pixels;<br>pixel size: 6.45 $\mu$m × 6.45 $\mu$m |

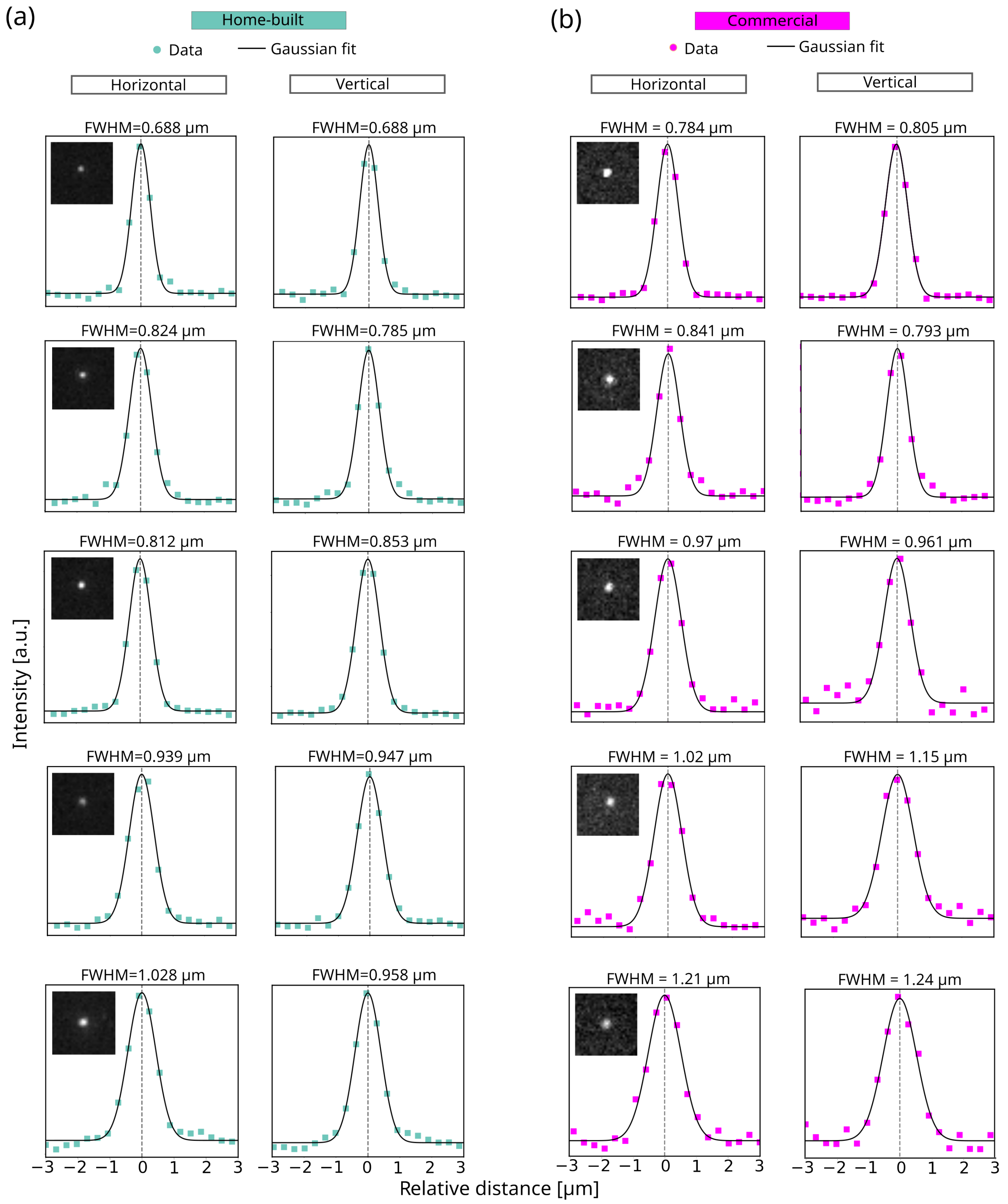


Fig. 1: Comparison of lateral point spread functions (PSFs) obtained using the home-built and commercial fluorescence microscopes. a) Representative PSFs from individual fluorescent beads imaged using the home-built (left, cyan) and b) the commercial (right, magenta) microscope setups. Each inset displays the corresponding bead image and the plots show line intensity profiles (colored circles) along the horizontal ($x$) and vertical ($y$) axes through the bead centroids with Gaussian fits (black solid lines).

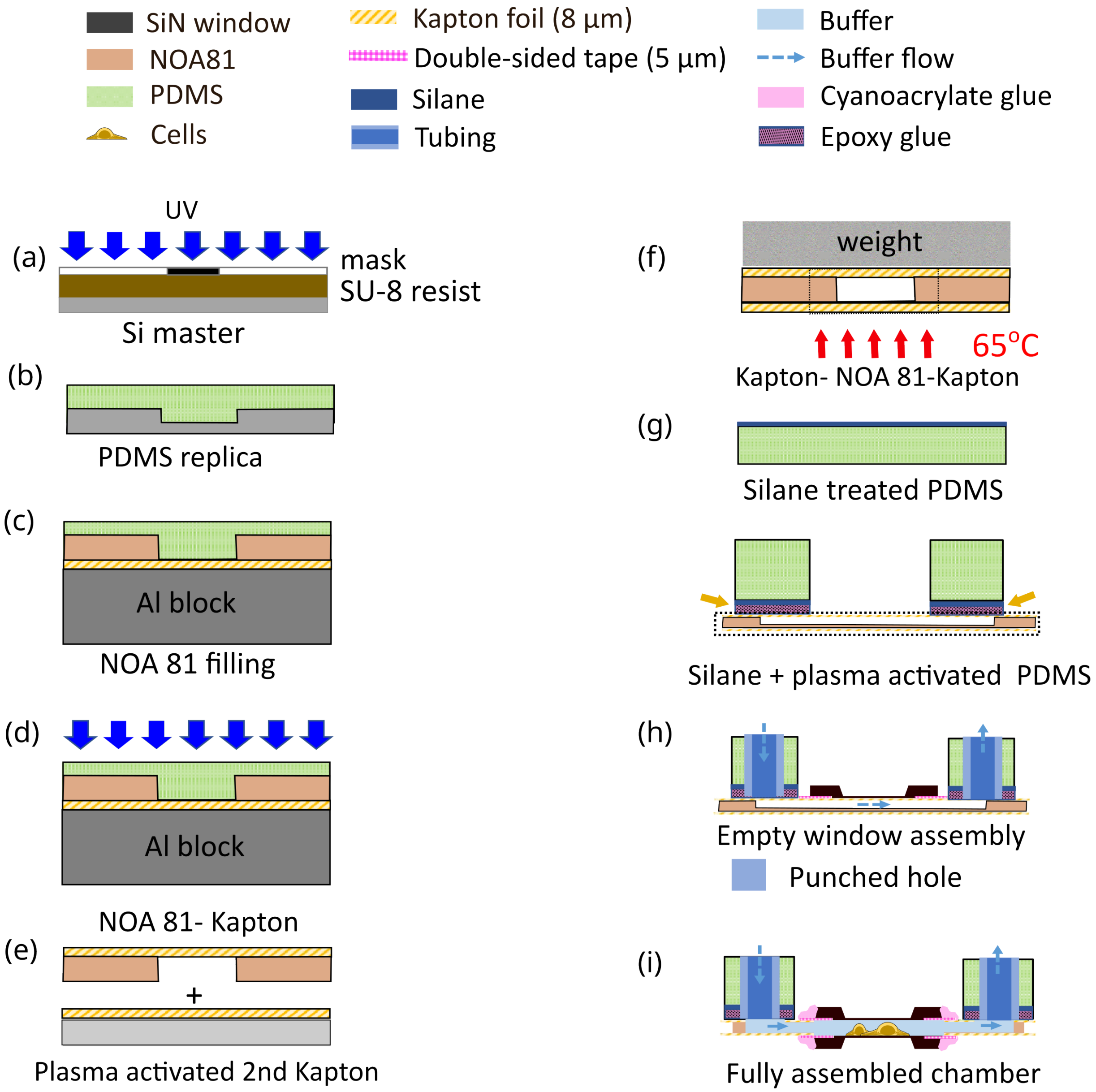


Fig. 2: Step-wise schematic of the fabrication workflow of the microfluidic chamber. (a) Photolithography to define the channel pattern by SU-8 photoresist on a silicon wafer. (b) PDMS replica molding from the silicon master. (c) NOA 81 replica molding on Kapton foil by capillary filling of the PDMS replica placed on an aluminum support and (d) UV curing of the NOA 81 forming a stable NOA 81-Kapton bond. (e) Sealing of the cured NOA 81-Kapton structure with a second, plasma activated Kapton foil. (f) Baking of the Kapton-NOA 81-Kapton structure (referred to as *device*) while being pressed down by a small weight. The dotted box indicates the channel region of the device. (g) Preparation of PDMS for device support. Bonding between the device and PDMS is achieved by silanization, plasma activation and applying epoxy glue. The activated side of PDMS support is indicated by orange arrows. (h) Punching holes in the center of device and the inlet/outlet. An empty $Si_3N_4$ window is attached to the center of the device with double-sided tape. (i) Fully assembled microfluidic chamber.

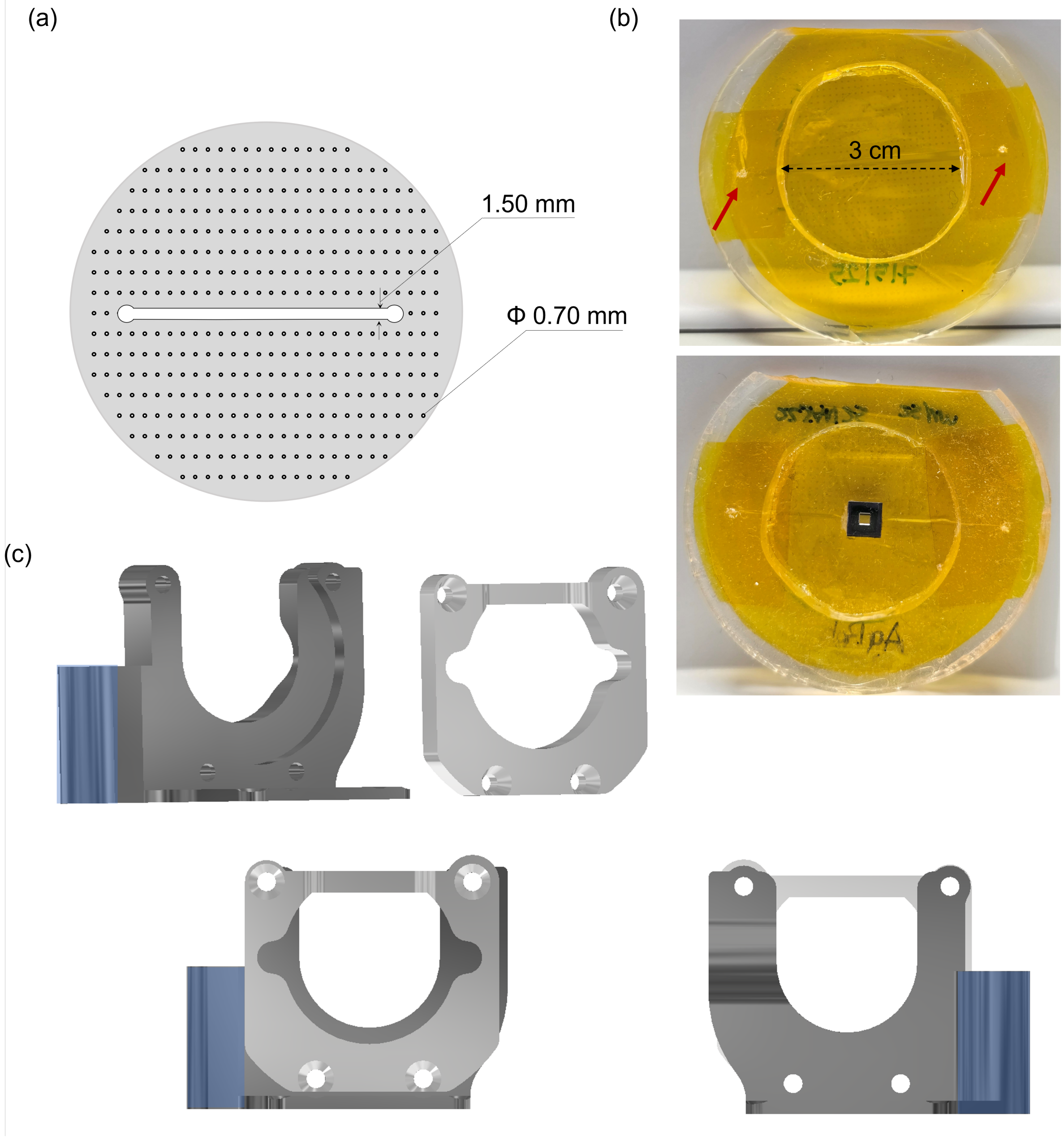


Fig. 3: (a) Photomask design. The outer diameter is 2 inches. The channel structure is centered on the mask; around the channel structure, we add periodic cylinders with a diameter of 700 $\mu$m and 1000 $\mu$m center-to-center distance as support structures to facilitate capillary filling (Supplementary Fig. 2c). (b) (Top) Photograph of the Kapton–NOA–Kapton device assembled on a PDMS support, forming the base of the microfluidic flow chamber. The inlet and outlet are indicated by red arrows. (Bottom) Image of the device after attaching an empty $Si_3N_4$ window. (c) CAD drawings of the chamber holder. (Top) Separate drawings of the two components. (Bottom) Front and back views of the fully assembled holder. The front faces the inlet and outlet tubing side of the ring–shaped PDMS support and is oriented towards the objective of the fluorescence microscope. The back view faces the incoming X-ray beam. The reservoir for collecting buffer from the outlet side is shown in blue.

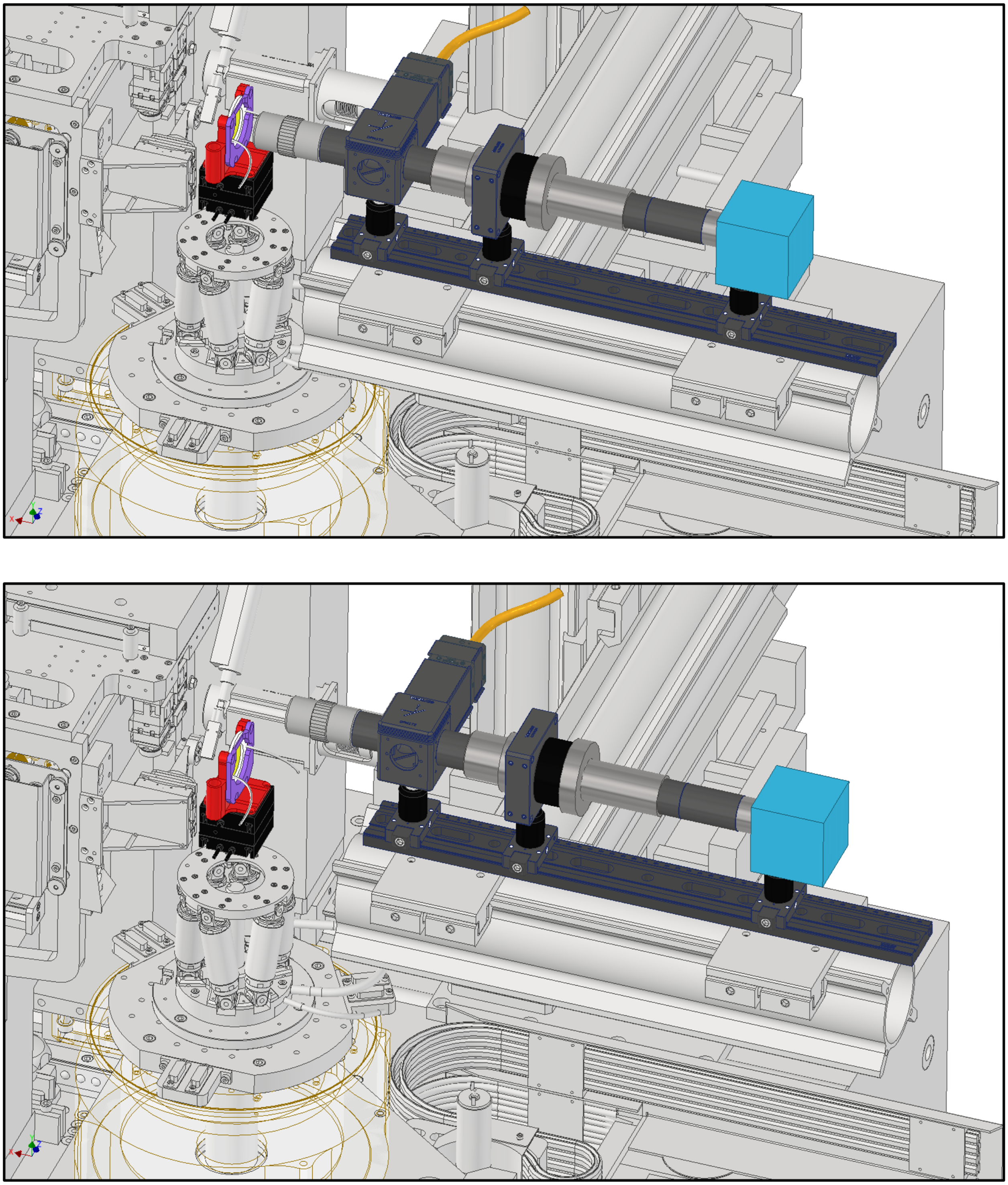

Fig. 4: CAD drawings of the fluorescence microscope set-up. (Top) Moved into the X-ray beampath and, therefore, aligned with the microfluidics chamber for the collection of fluorescence images of the sample. (Bottom) Moved out of the X-ray beampath. The translation range is 125 cm.

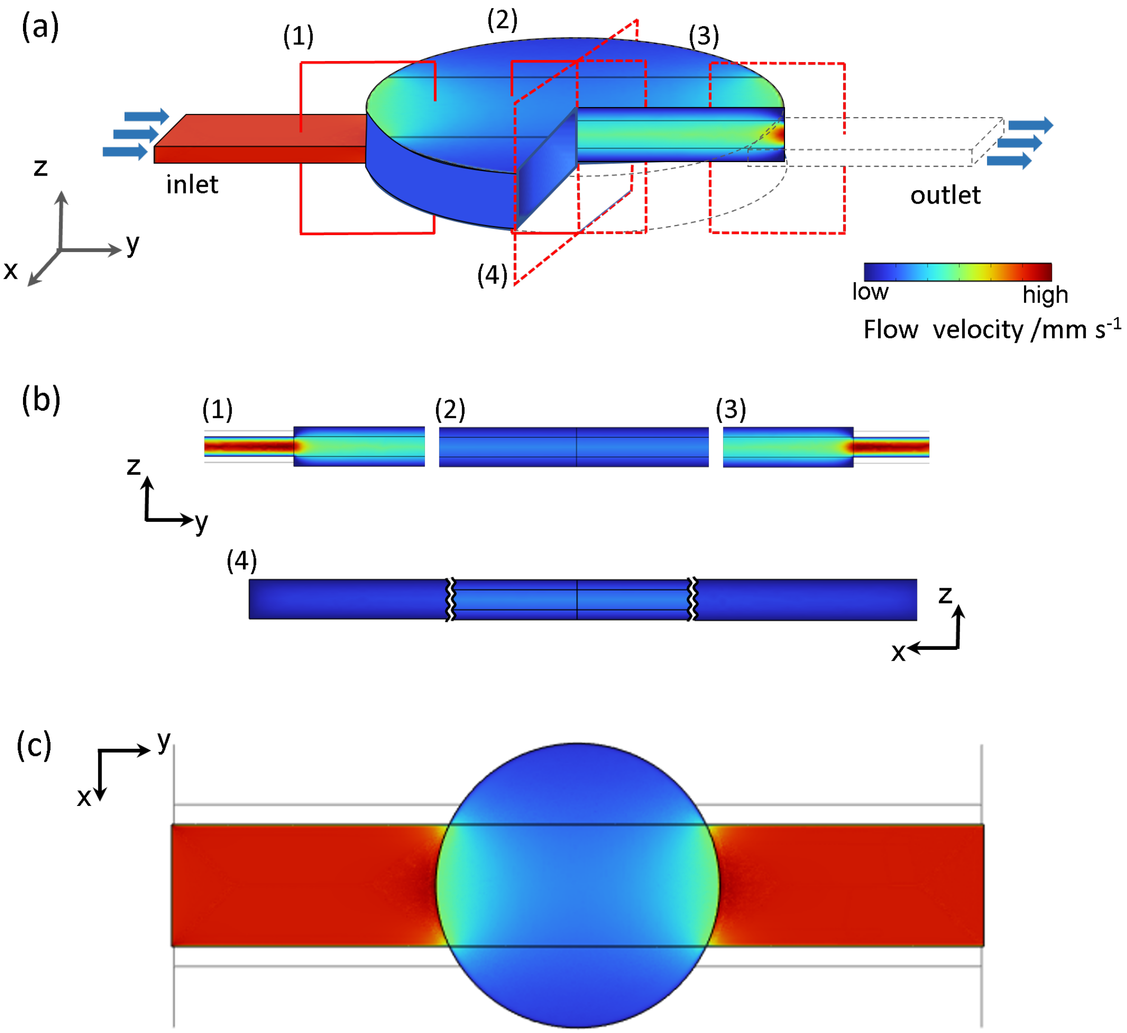


Fig. 5: Finite element method (FEM) simulation of the flow conditions within the microfluidic chamber (COMSOL multiphysics 6.2). (a) 3D representation of the flow velocity in mm/s within the straight microchannel and observation region in the center defined by a "sandwich" of two $Si_3N_4$ windows. Arrows indicate the flow direction. The color map represents the simulated flow velocity, highlighting reduced flow velocity inside the observation region compared to the narrow channels. (b) Longitudinal cross-sections showing the evolution of the flow velocity from inlet to outlet for four specific regions indicated as (1) to (4) in panel a. Regions (2) and (4) correspond to the membrane region of the $Si_3N_4$ window, where fluorescence microscopy and scanning SAXS measurements are performed. (c) Top view illustrating the spatial distribution of flow velocities across the observation region–channel interface.

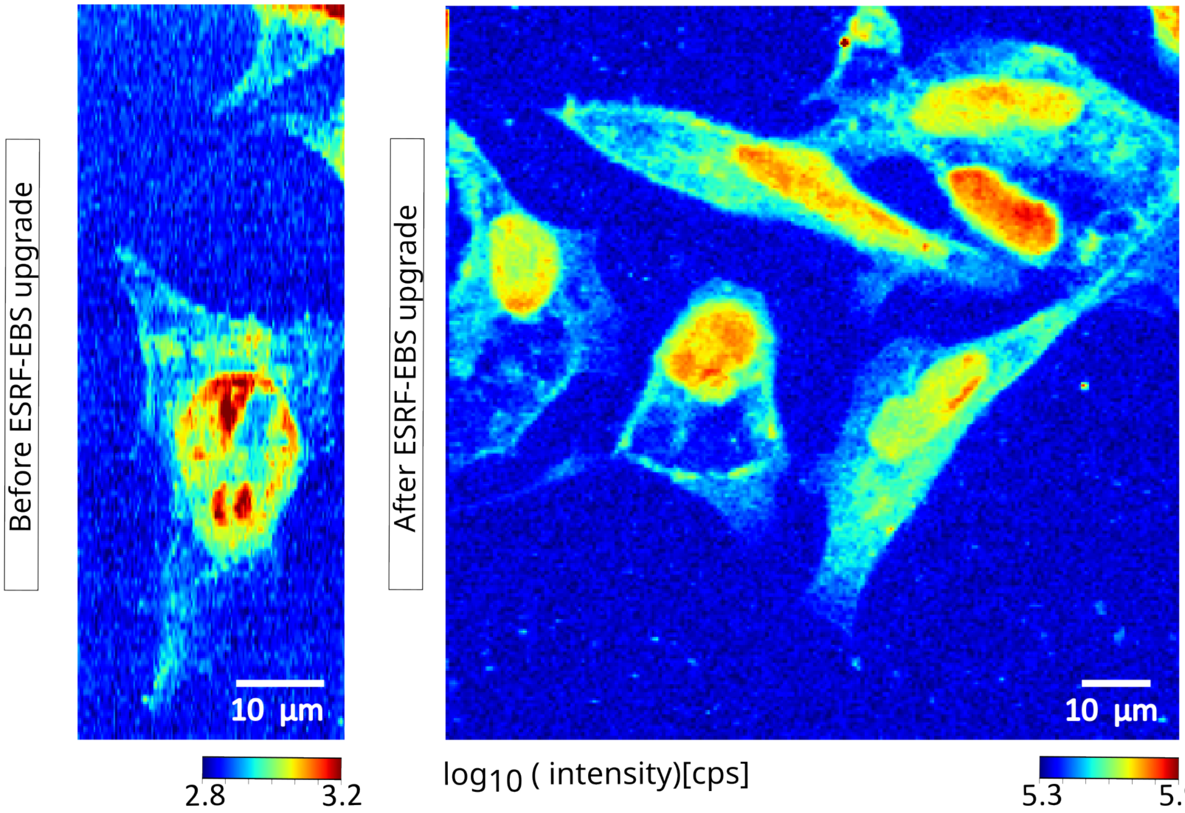


Fig. 6: X-ray dark-field images of fixed-hydrated SW13 cells before and after the "Extremely Brilliant Source" (EBS) upgrade at ESRF. The water layer thickness of the microfluidic chamber before the upgrade was 200 $\mu$m, whereas the newly designed microfluidic chamber compatible with both optical imaging and fast scanning SAXS modalities has a water layer thickness of 60$\pm$ 10 $\mu$m. The scanning parameters for the respective measurements are listed in Supplementary Table 2. The $q$-range considered for calculating the data before the upgrade is (0.175–0.262) nm$^{-1}$, whereas for the present beamtime it is (0.19–0.39) nm$^{-1}$

Table 2: Parameters for fast scanning SAXS before and after the EBS upgrade at ESRF

| **Parameters** | **Pre upgrade** | **Post upgrade** |
|---|---|---|
| Photon energy [keV] | 14.92 | 15.00 |
| Photon flux [ph/s] | $8.7 \times 10^9$ | $1.2 \times 10^{12}$ |
| Exposure time [s] | 0.5 | 0.002 |
| Step size ($\Delta_{y,z}$ [$\mu$m$^2$]) | $0.3 \times 1$ | $0.5 \times 0.5$ |
| Dose per scan point [Gy] | $5.3 \times 10^6$ | $6.3 \times 10^6$ |
| Scattering contrast $\left(\frac{I_{\text{cell}}-I_{\text{bkg}}}{I_{\text{bkg}}}\right)$ | 1.5 | 3.0 |

Table 3: Calculated flow velocities in the channel and the observation region for different flow rates

| **Location** | **Width ($\mu$m)** | **Height ($\mu$m)** | **Cross-sectional area ($\mathbf{m^2}$)** | **Flow rate ($\mu$L/h)** | **Flow velocity (mm/s)** |
|---|---|---|---|---|---|
| Observation region | 3500 | 60 | $2.1 \times 10^{-7}$ | 20 | 0.026 |
| | | | | 50 | 0.063 |
| | | | | 100 | 0.132 |
| | | | | 200 | 0.263 |
| Channel | 1500 | 25 | $2.25 \times 10^{-8}$ | 20 | 0.246 |
| | | | | 50 | 0.62 |
| | | | | 100 | 1.24 |
| | | | | 200 | 2.48 |

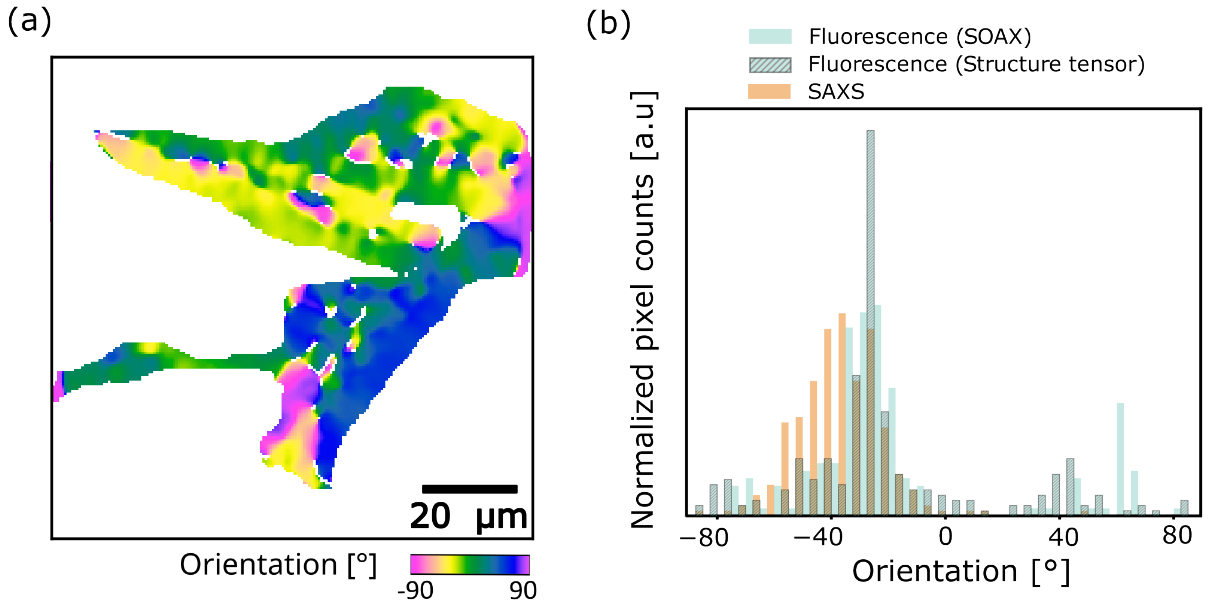


Fig. 7: (a) Orientation map derived from the fluorescence image of keratin networks in SW-13 cells (see Fig. 3a in the main text) using the structure tensor method. (b) Comparison of the orientation histograms obtained from SAXS and fluorescence analysis using SOAX (as shown in the main text) and the structure tensor method. Only the pixels within the white outlined region of Fig. 3d of the main text are included in this analysis.

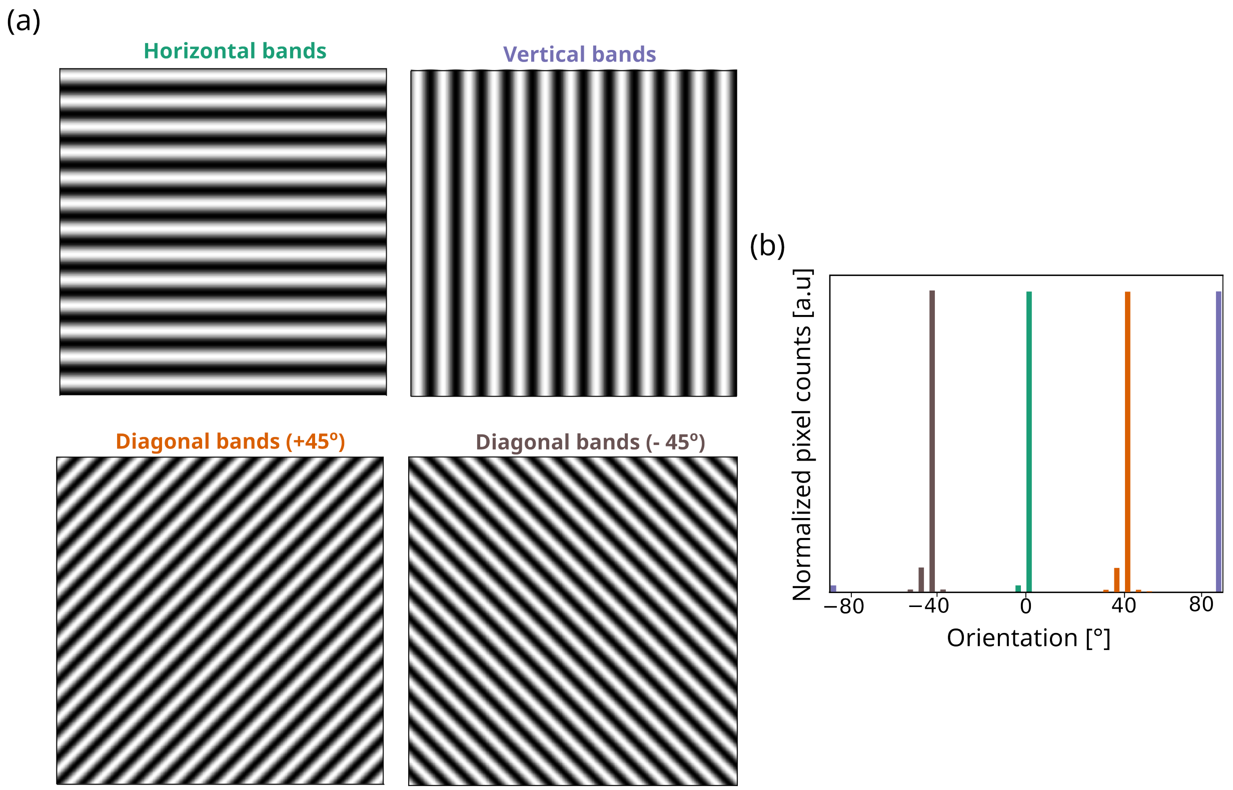


Fig. 8: Application of the structure tensor method to synthetic data. (a) Grayscale images containing horizontal, vertical and diagonal (+45° and -45°) stripe patterns. (b) The corresponding orientation histograms show distinct peaks at the expected angles, demonstrating that the structure tensor analysis reliably captures dominant alignment across different orientations.
